# Performance Assessment and Selection of Normalization Procedures for Single-Cell RNA-seq

**DOI:** 10.1101/235382

**Authors:** Michael B. Cole, Davide Risso, Allon Wagner, David DeTomaso, John Ngai, Elizabeth Purdom, Sandrine Dudoit, Nir Yosef

**Affiliations:** Department of Physics, University of California, Berkeley; Division of Biostatistics and Epidemiology, Department of Healthcare Policy and Research, Weill Cornell Medicine, New York, NY; Department of Electrical Engineering and Computer Sciences, University of California, Berkeley; Center for Computational Biology, University of California, Berkeley; Department of Molecular and Cell Biology, University of California, Berkeley; Department of Statistics, University of California, Berkeley; Division of Epidemiology and Biostatistics, School of Public Health, University of California, Berkeley

## Abstract

Systematic measurement biases make data normalization an essential preprocessing step in single-cell RNA sequencing (scRNA-seq) analysis. There may be multiple, competing considerations behind the assessment of normalization performance, some of them study-specific. Because normalization can have a large impact on downstream results (e.g., clustering and differential expression), it is critically important that practitioners assess the performance of competing methods.

We have developed *scone* — a flexible framework for assessing normalization performance based on a comprehensive panel of data-driven metrics. Through graphical summaries and quantitative reports, *scone* summarizes performance trade-offs and ranks large numbers of normalization methods by aggregate panel performance. The method is implemented in the open-source Bioconductor R software package scone. We demonstrate the effectiveness of *scone* on a collection of scRNA-seq datasets, generated with different protocols, including Fluidigm C1 and 10x platforms. We show that top-performing normalization methods lead to better agreement with independent validation data.

## Introduction

Normalization is a common preprocessing step in the analysis of -omic data, such as high-throughput transcriptome microarray and sequencing (RNA-seq) data. The goal of normalization is to account for observed differences in measurements between samples and/or features (e.g., genes) resulting from technical artifacts or unwanted biological effects (e.g., batch effects) rather than biological effects of interest. Accordingly, two types of normalization are often considered: between-sample and within-sample. This article focuses on the former. In order to derive gene expression measures from single-cell RNA sequencing (scRNA-seq) data and subsequently compare these measures between cells, analysts must normalize read counts (or other expression measures) to adjust for obvious differences in sequencing depths. When there are other significant biases in expression quantification, it may be necessary to further adjust expression measures for more complex unwanted technical factors related to sample and library preparation.

As previously discussed [1, 2], normalization of scRNA-seq data is often accomplished via methods developed for bulk RNA-seq or even microarray data. These methods tend to neglect prominent features of scRNA-seq data such as: *zero inflation*, i.e., an artifactual excess of zero read counts observed in some single-cell protocols (e.g., SMART-seq) [3, 4]; transcriptome-wide nuisance effects (e.g., batch), comparable in magnitude to the biological effects of interest [5]; uneven sample quality, e.g., in terms of alignment rates and nucleotide composition [6]. In particular, widely-used global-scaling methods, such as reads per million (RPM) [7], trimmed mean of M values (TMM) [8], and DESeq [9], are not well suited to handle large or complex batch effects and may be biased by low counts and zero inflation [2]. Other more flexible methods, such as remove unwanted variation (RUV) [10, 11] and surrogate variable analysis (SVA) [12, 13], depend on tuning parameters (e.g., the number of unknown factors of unwanted variation).

A handful of normalization methods specifically designed for scRNA-seq data have been proposed. These include scaling methods [14, 15], regression-based methods for known nuisance factors [16, 17], and methods that rely on spike-in sequences from the External RNA Controls Consortium (ERCC) [18, 19]. While these methods address some of the problems affecting bulk normalization methods, each suffers from limitations with respect to their applicability across diverse study designs and experimental protocols. Global-scaling methods define a single normalization factor per cell and thus are unable to account for complex batch effects. Explicit regression on known nuisance factors (e.g., batch, number of reads in a library) may miss unknown, yet unwanted variation, which may still confound the data [11]. Unsupervised normalization methods that regress gene expression measures on unknown unwanted factors may perform poorly with default parameters (e.g., number of factors adjusted for) and require tuning, while ERCC-based methods suffer from differences between endogenous and spiked-in transcripts [2, 11]. Protocols using unique molecular identifiers (UMI) still require normalization; while UMIs remove amplification biases, they are often sensitive to sequencing depth and differences in capture efficiency before reverse transcription [2].

Due to the prevalence of confounding in single-cell experiments, the lack of a uniformly optimal normalization across datasets, and the ambiguity in tuning parameter guidelines for commonly-used normalization methods, we recommend the inspection and evaluation of many approaches and the use of multiple data-driven metrics to guide the selection of suitable approaches for a given dataset. We have developed the *scone* framework for implementing and assessing the performance of a range of *normalization procedures*, each consisting of defined normalization steps, such as scaling and supervised or unsupervised regression-based adjustments. *scone* evaluates the performance of each procedure and ranks them by aggregating over a panel of performance metrics that consider different aspects of a desired normalization outcome, including both removal of unwanted variation and preservation of wanted variation.

We demonstrate that the *scone* methodology is generally applicable to different scRNA-seq protocols and study designs. The modularity of the Bioconductor R software package scone allows researchers to tune and compare a set of default normalizations as well as to include user-defined methods, providing a useful framework for both practitioners and method developers.

## Results

### scRNA-seq data are affected by batch effects and other unwanted variation

We briefly illustrate the difficulties of normalizing single-cell RNA-seq data with a published SMART-seq dataset, processed on Fluidigm C1 [20]. We focus on a subset of 420 mouse Th17 T-cells harvested after *in vitro* differentiation of CD4+ naive T-cells under 48 hours of pathogenic (IL-1*β*+IL-6+IL-23) or non-pathogenic (TGF-*β*1+IL-6) conditioning. Prior to conditioning, these cells had been extracted from two strains of mice: wild-type B6 and transgenic B6 mice with an IL17a GFP reporter. We have aligned the publicly available reads to the mouse genome, counting over RefSeq gene intervals to generate a cells-by-genes count matrix. After sample and gene filtering, 337 libraries and 7,590 genes are retained for normalization and downstream analysis, preserving ~ 80% of all reads and cells (Supplementary Fig.1b; see Methods).

Even after scaling the gene-level read counts by total counts (TC), mouse-specific effects are prominently featured in *principal component analysis* (PCA; Fig. 1a). In the space defined by the first two *principal components* (PC), the distances between cells from mouse 7 and mouse 8 are larger than the distances between pathogenic and non-pathogenic cells collected from the same mouse. Due to the partially confounded design, these mouse effects may result from multiple sources, including (i) true biological differences between mice or (ii) mouse-specific technical biases. The study design prevents us from teasing apart these two effects, but we can account for technical contributions to the read counts by examining the association of the expression PCs with RNA-seq library QC metrics (Fig. 1b; see Tables 1 and 2 and Methods).

**Figure 1:**
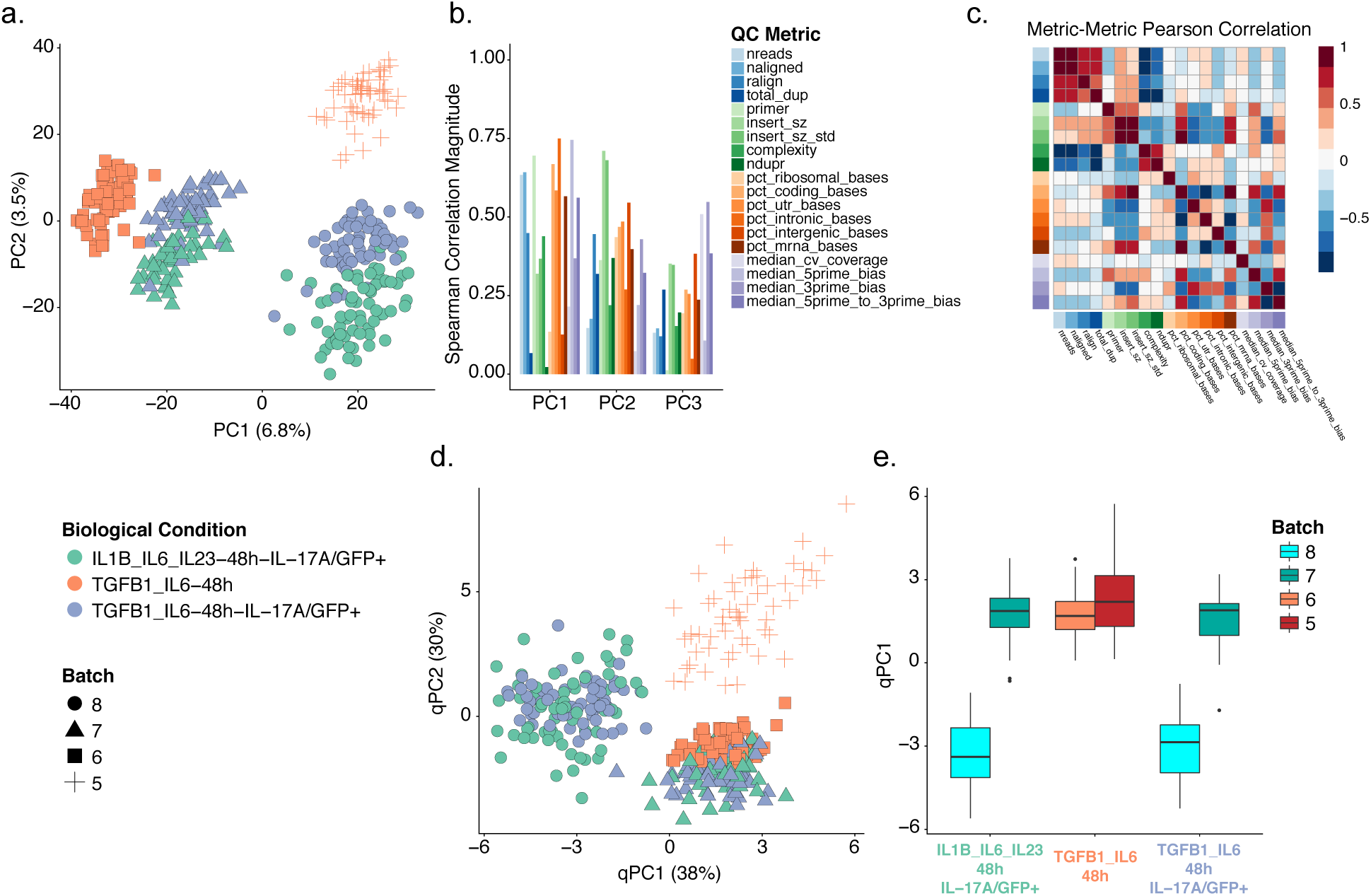
Exploratory data analysis of mouse Th17 dataset [20]. (a) Principal component analysis (PCA) of the log-transformed, total-count-normalized (TC) read count data for all genes and cells passing quality filtering (see Methods). Cells are color-coded by biological condition; shape represents the donor mouse (batch). For two of the three conditions, samples were extracted from only one mouse (IL1B_IL6_IL23-48h-IL-17A/GFP+ and TGFB1_IL6–48h-IL-17A/GFP+ from mice 7 and 8, respectively), while samples from the third condition (TGFB1_IL6–48h) came from two distinct mice (mice 5 and 6). Cells cluster by both biological condition and batch, the latter representing unwanted variation. (b) Absolute Spearman correlation coefficient between the first three principal components (PCs) of the expression measures (as computed in (a)) and a set of quality control (QC) measures (Table 1). (c) Heatmap of pairwise Pearson correlation coefficients between QC measures. (d) PCA of the QC measures for all cells in (a). PCs of QC measures are labeled “qPCs” to distinguish them from expression PCs. Single-cell QC profiles cluster by batch, representing important aspects of batch covariation. (e) Boxplot of the first qPC, stratified by both biological condition and batch. Note that there are different numbers of cells in each stratum.

**Table 1:** Sample-level quality control (QC) measures (non-10x Genomics).

| Name | Description | Source |
| --- | --- | --- |
| NREADS | Total number of sequenced reads | Picard |
| NALIGNED | Total number of aligned reads | Picard |
| RALIGN | Percentage of mapped reads | Picard |
| TOTAL_DUP | Number of duplicate reads | FastQC |
| PRIMER | Percentage of primer sequence reads | FastQC |
| INSERT_SZ | Average insert size | Picard |
| INSERT_SZ_STD | Insert size variance | Picard |
| COMPLEXITY | Sequence Complexity | Picard |
| NDUPR | Percentage of unique reads | Picard |
| PCT_RIBOSOMAL_BASES | Percentage of ribosomal bases | Picard |
| PCT_CODING_BASES | Percentage of coding bases | Picard |
| PCT_UTR_BASES | Percentage of UTR bases | Picard |
| PCT_INTRONIC_BASES | Percentage of intronic bases | Picard |
| PCT_INTERGENIC_BASES | Percentage of intergenic bases | Picard |
| PCT_MRNA_BASES | Percentage of mRNA bases | Picard |
| MEDIAN_CV_COVERAGE | Median coefficient of variation of coverage | Picard |
| MEDIAN_5PRIME_BIAS | Mean 5' coverage bias | Picard |
| MEDIAN_3PRIME_BIAS | Mean 3' coverage bias | Picard |
| MEDIAN_5PRIME_TO_3PRIME_BIAS | Mean 5' to 3' coverage bias | Picard |

**Table 2:** Sample-level quality control (QC) measures (10x Genomics).

| Name | Description | Source |
| --- | --- | --- |
| num_umi | Number of unique UMI sequences | Cell Ranger |
| num_reads | Total number of reads (regardless of mapping) | Cell Ranger |
| mean_reads_per_umi | The average number of reads supporting each UMI | Cell Ranger |
| std_reads_per_umi | Standard deviation of the number of reads supporting each UMI | Cell Ranger |
| mapped_reads | Proportion of reads which confidently mapped to a gene | Cell Ranger |
| genome_not_gene | Proportion of reads mapping to the genome, but not to a gene | Cell Ranger |
| unmapped_reads | Proportion of reads which did not align | Cell Ranger |
| umi_corrected | Proportion of reads whose UMI sequence was corrected by Cell Ranger | Cell Ranger |
| barcode_corrected | Proportion of reads whose barcode sequence was corrected by Cell Ranger | Cell Ranger |

The first three expression PCs exhibit large correlations with measures of genomic alignment rate, primer contamination, intronic alignment rate, and 5’ bias (see Methods). The correlation structure between these QC measures reflects constraints on library quality in the study (Fig. 1c). While some of these pairwise associations represent natural dependencies between similar QC measures (e.g., total number of reads and total number of aligned reads are positively correlated), others reflect mouse-specific technical biases in the study. Applying PCA to the matrix of QC measures, we can see how these metrics provide a candidate basis for representing batch (i.e., mouse) effects (Fig. 1d). Inter-batch QC differences are relatively large as in Figure 1a, while intra-batch differences between pathogenic and non-pathogenic cells are noticeably smaller (Fig. 1e). Mouse 6 libraries are technically similar to mouse 7 libraries, while cells from mice 5 and 8 are technically distinct; these relationships are similar to those observed in the PCA of gene expression measures, suggesting that the corresponding structure in the expression data is artifactual. It is surely possible that some of the observed associations between read counts and QC measures result from biological confounding rather than direct technical bias, i.e., a cell’s biological state may impact transcriptome integrity and sequencing viability [2, 21]. However, unlike factors such as mouse-of-origin, there exist simple interpretations for correlations between quantified expression measures and library alignment statistics.

We have focused on a specific dataset in this section, but we note that many of these observations are not unique to this example and are general features of scRNA-seq. To highlight this, we have performed similar exploratory data analyses on a set of developing human cortical neurons assayed using a 2014 Fluidigm protocol [22] and a set of human peripheral blood mononuclear cells (PBMC) assayed using the 10x Chromium platform [23] (Supplementary Fig. 2 and 3).

### *scone*: An exploratory framework for the implementation and evaluation of scRNA-seq normalization

As illustrated in Figure 1, simple global scaling alone is insufficient for normalizing scRNA-seq data and more flexible and aggressive procedures aimed at removing unwanted variation (e.g., batch effects) may be generally beneficial. Here, we present a general framework for implementing and evaluating normalization procedures for scRNA-seq data (Fig. 2).

**Figure 2:**
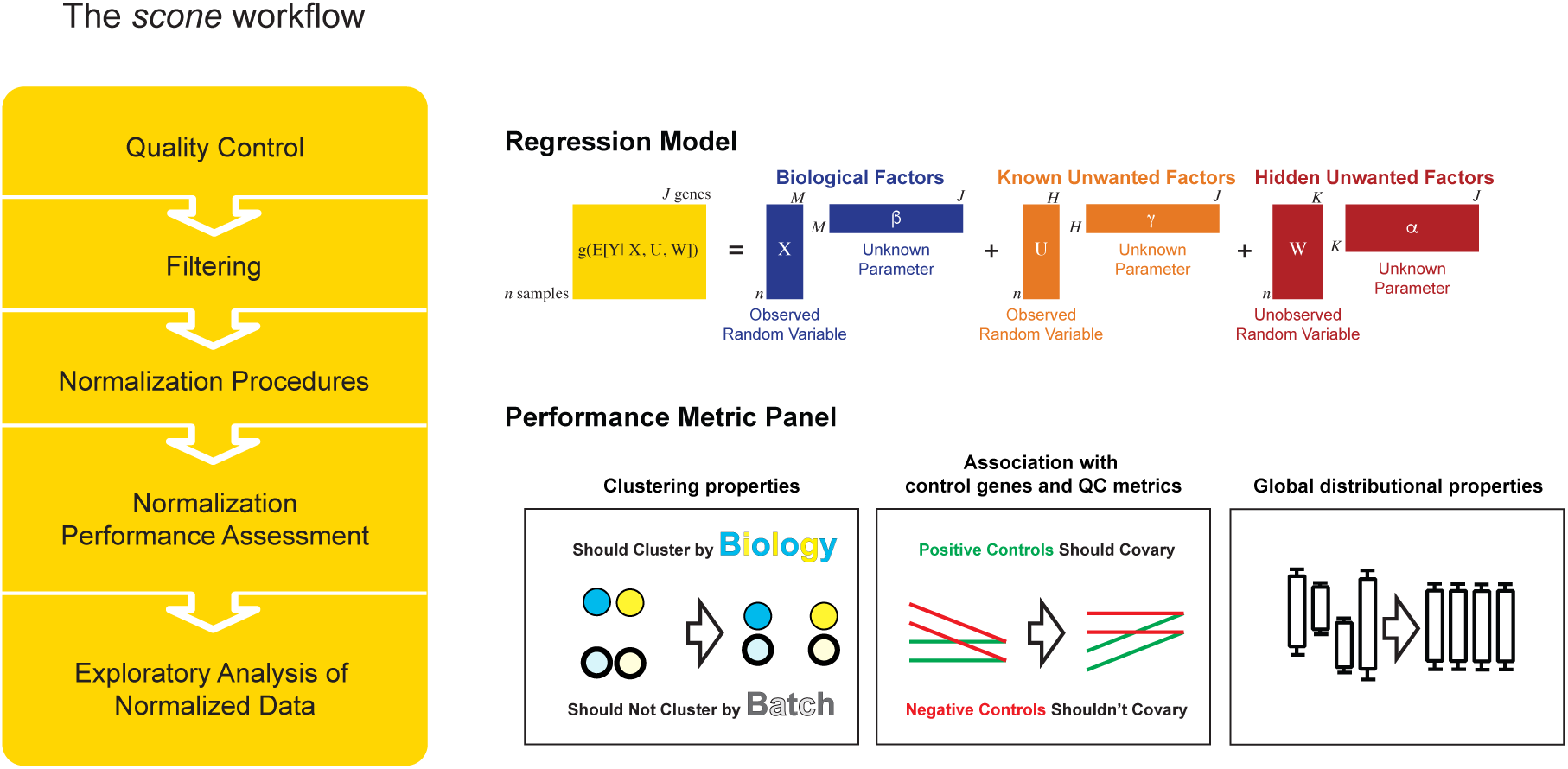
Schematic view of the *scone* workflow. The yellow box summarizes the five main steps of the *scone* workflow. (i) QC measures are obtained for each sample, using Picard tools (Table 1), Cell Ranger (Table 2), or other tools such as scater. (ii) (Optional) Sample-level QC measures are used to filter out low-quality samples. Subsequently, lowly-expressed genes are identified and filtered out to reduce the impact of noisy features on downstream analysis. (iii) Data are normalized via many combinations of scaling procedures and regression-based procedures, modeling both known and unknown variation as indicated in the regression model diagram. (iv) Normalized data are evaluated and ranked according to a panel of eight performance metrics, spanning three categories: (a) Clustering properties (e.g., removing batch effects and preserving biological heterogeneity), (b) association with control genes and QC metrics (e.g., preserving association with positive controls and removing association with QC measures), and (iii) global distributional properties (e.g., reducing global expression variability). (v) One or more highly-ranked normalization procedures are analyzed in parallel and downstream conclusions are compared.

The full *scone* pipeline consists of two steps prior to normalization, namely (i) QC assessment and (ii) optional sample and gene filtering, resulting in the selection of high-quality cell gene expression profiles.

Following these initial steps, *scone* uses a two-part normalization template to define an ensemble of normalization procedures: (i) scaling of the counts to account for between-sample differences in sequencing depth and other parameters of the read count distributions; (ii) regression-based adjustment for known unwanted factors, such as processing batches, and unknown unwanted factors [11]. Figure 2 displays some of the methods that can be employed in our normalization template. For instance, one can apply either TC scaling or more robust scaling procedures designed to reduce the effect of outliers (e.g., TMM [8] or DESeq [9]). Additionally, confounding factors can be adjusted for by regressing scaled gene expression measures on quantities known to influence them (e.g., batches or the QC measures shown in Figure 1). Alternatively, unsupervised procedures can estimate hidden unwanted factors and regress them out of the data (e.g., RUV of Risso *et al*. [11]).

The evaluation step of *scone* involves the comparison and ranking of all normalization procedures to identify sets of top performing procedures. To achieve this, *scone* calculates a set of eight performance metrics, aimed at capturing different aspects of successfully normalized data. We classify these metrics into three broad categories: (i) clustering of samples according to factors of wanted and unwanted variation; (ii) association of expression principal components with factors of wanted and unwanted variation; (iii) between-sample distributional properties of the expression measures (see Methods). Overall, these metrics capture the trade-offs between the ability of a normalization procedure to remove unwanted variation, preserve biological variation of interest, and maintain minimum global technical expression variability. These trade-offs are rooted in the confounding commonly encountered in single-cell assays; *scone* provides a reproducible basis for managing those trade-offs via normalization. A simple ranking of normalization procedures can be obtained by averaging ranks based on the eight individual metrics, although the multidimensional aspect of normalization performance is lost in this one-dimensional representation. To account for this, we advocate the inspection of the full space of normalization performance measures, through specialized plots (Fig. 3a) and the *scone* report browser (Fig. 8). Notably, in order to avoid an evaluation dominated by how normalization procedures handle zeros, we force to zero all values that are initially zeros as well as any negative values produced by a normalization procedure (see Methods).

**Figure 3:**
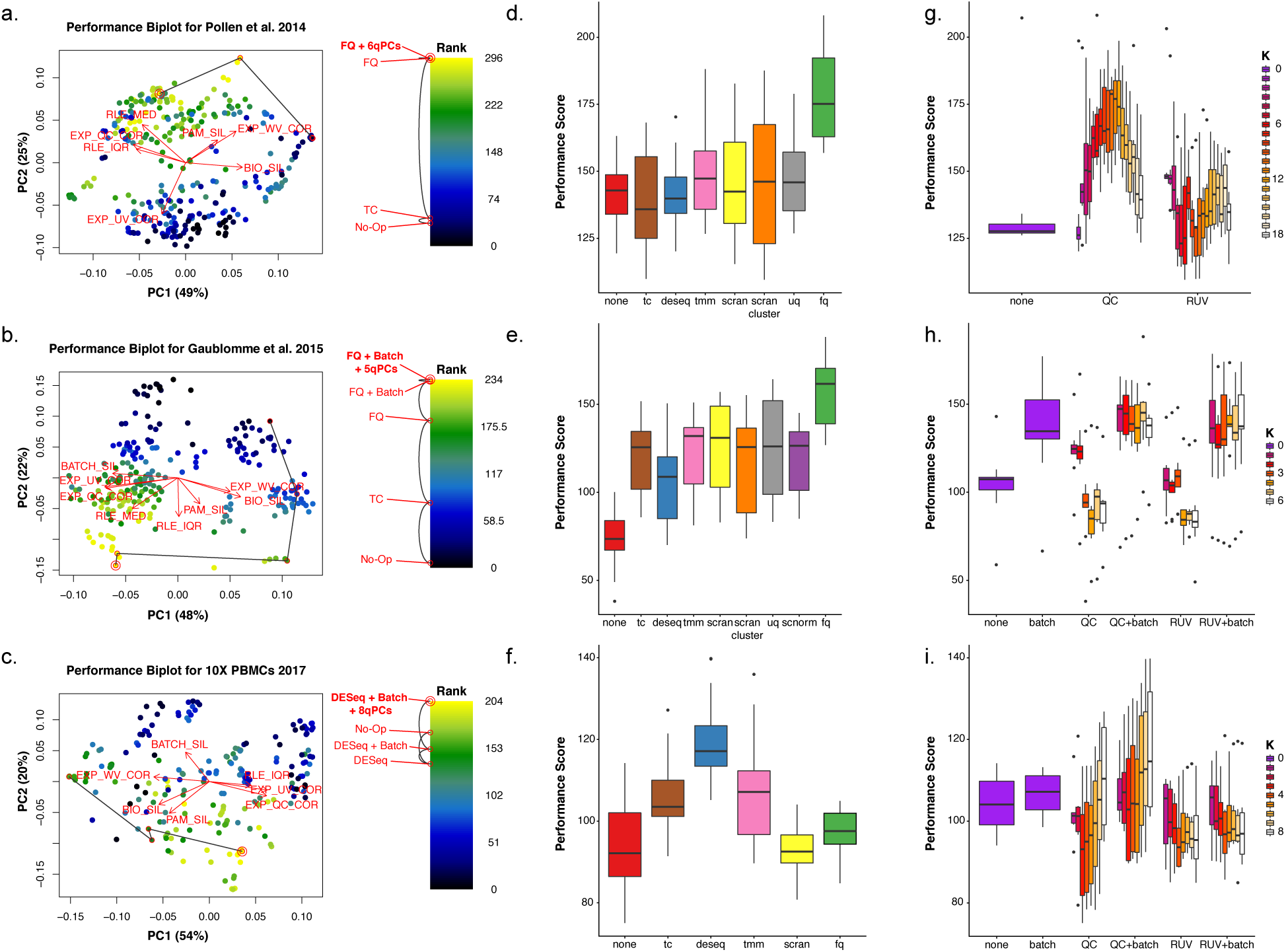
Normalization performance assessment for three scRNA-seq datasets [20, 22, 23]. (a-c) Biplot [24] showing the first two PCs of eight rank-transformed *scone* performance metrics, or fewer if some are undefined or invariant: Preservation of biological clustering (“BIO_SIL”), batch effect removal (“BATCH_SIL”), cluster heterogeneity (“PAM_SIL”), preservation of association with positive control genes (“EXP_WV_COR”), removal of unwanted associations (negative control genes, “EXP_UV_COR”, or sample-level QC measures, “EXP_WC_COR”), and global distributional uniformity (“RLE_MED” and “RLE_IQR”). Each point corresponds to a normalization procedure and is color-coded by the rank of the *scone* performance score (mean of eight *scone* performance metric ranks). The red arrows correspond to the PCA loadings for the eight performance metric ranks. The direction and length of a red arrow can be interpreted as a measure of how much each metric contributes to the first two PCs. Red circles mark the best normalization (w/ double circle), no normalization, and other normalization procedures relating the two (see labels). Key: ‘No-Op” = No normalization, “TC” = Total-count normalization, “FQ” = Full-quantile normalization, “DESeq” = Relative log-expression scaling [9], “Batch” = Regression-based batch normalization, “*k*qPCs” = Regression-based adjustment for first *k* qPCs. (d-f) Boxplot of *scone* performance score, stratified by scaling normalization method, for the three scRNA-seq datasets presented in the same order as in (a-c). (g-i) Boxplot of *scone* performance score, stratified by regression-based normalization method (batch, QC, and RUV), for the three scRNA-seq datasets presented in the same order as in (a-c).

Below we draw on evidence from multiple single-cell datasets, generated from various technological platforms, to show how no single normalization method is uniformly optimal: insights from the *scone* framework will highlight how performance depends on the design of the experiment and other characteristics of the data.

### *scone* removes unwanted variation while preserving biological variation of interest

A useful representation of the normalization performance landscape is the *biplot* [24], in which each point corresponds to a normalization procedure and the dimensions of variation, represented by red arrows, correspond to *scone* performance metrics (Fig. 3a-c).

The *scone* biplot naturally represents trade-offs between these metrics, as illustrated in Figure 3 for three datasets. For the cortical neuron dataset of Pollen *et al*. [22] (Fig. 3a), there are two major bundles of red arrows, representing (i) preservation of biological heterogeneity and (ii) distributional uniformity irrespective of library quality. The existence of this trade-off suggests that wanted variation is confounded with measurement artifacts. The top-ranked procedure according to *scone* involved full-quantile (FQ) normalization followed by adjustment for 6 QC PCs. Tracing a path from the performance coordinates of unnormalized data to those of the top-ranked normalization, we notice that FQ without adjustment also performs very well according to *scone*, occupying a middle position between these two trade-offs. Both procedures may reasonably be carried to downstream analysis, but the *scone* biplot highlights an important tension in this decision.

The *scone* biplot for the SMART-seq dataset of Gaublomme *et al*. [20] demonstrates a more complex “fan” of trade-offs between batch effect removal and preservation of wanted variation (Fig. 3b). Compared to no normalization, global-scaling and full-quantile normalization primarily improve distributional properties of the data, reducing the amount of global expression variability between samples (captured by the RLE metrics; see Methods). Regression-based normalization, including batch regression and RUV, remove unwanted variation at the expense of wanted variation; the biplot can help identifying those normalization that balance the trade-off between removing too much biological variation and too little technical variation. Unlike the biplot for Pollen *et al*. [22], the arrows corresponding to associations with QC metrics or negative control genes are closely aligned. However, the factors of unwanted variation computed from negative control genes (via RUVg) and the QC measures are not always correlated (Supplementary Fig. 4), suggesting that regression normalizations based on these factors are complementary approaches.

We observe similar trade-offs between removal of technical variation and preservation of biological variation in the 10x Genomics dataset (Fig. 3c). Unique to this case is the relatively good performance of no normalization: most normalization procedures scored worse than doing nothing. Nevertheless, *scone* identifies a procedure that balances the trade-offs between the different metrics, involving the DESeq scaling factor and regression-based adjustment for all 8 PCs of the QC matrix.

### *scone’s* normalization performance ranking is data-adaptive

Global-scaling normalization methods are ranked similarly for both C1 datasets (Fig. 3d-e), under-performing the more aggressive full-quantile normalization. Importantly, for the dataset of Pollen et al., globally-scaled data do not show improved performance when compared to unscaled data (Fig. 3b). Conversely, scaling by *DESeq* size factors outperforms other scaling normalizations as well as full-quantile normalization for the 10x Genomics PBMC dataset (Fig. 3f). For all of these datasets, single-cell-specific methods, such as those implemented in the R packages scran and SCnorm, do not outperform methods developed for bulk RNA-seq.

The inclusion of a batch regression step in the normalization strategy is desirable for the SMART-seq dataset; procedures including QC or RUV factors without batch normalization perform poorly (Fig. 3h). This result indicates that, for this study, preexisting batch classifications are better proxies of inter-batch effects than QC or RUVg factors, despite their problematic associations with biological condition. In contrast, QC-based regression normalization outperforms RUVg for the Pollen et al. dataset (Fig. 3g), as well as in the 10x dataset when paired with batch adjustment (Fig. 3i). Taken together, these observations suggest that there is no single normalization method that uniformly outperforms the others and that *scone* is able to identify appropriate normalization procedures in a data-dependent fashion.

### *scone’s* subsample-based normalization performance ranking approximates full-data ranking

One potential drawback of *scone* is its computational complexity, implementing and ranking hundreds of normalization procedures per dataset. This can be especially problematic when applying *scone* to large datasets. In such cases, an efficient strategy is to use a random subset of the cells for the purpose of ranking normalizations, applying only the best performing normalization procedure to the full dataset. For the 10x PBMC dataset, this subsampling strategy leads to a ranking that is highly consistent with the ranking based on the full data, as illustrated in Figure 4. Importantly, as little as 10% of the cells is enough to yield more than 80% correlation with the full ranking (Fig. 4d).

**Figure 4:**
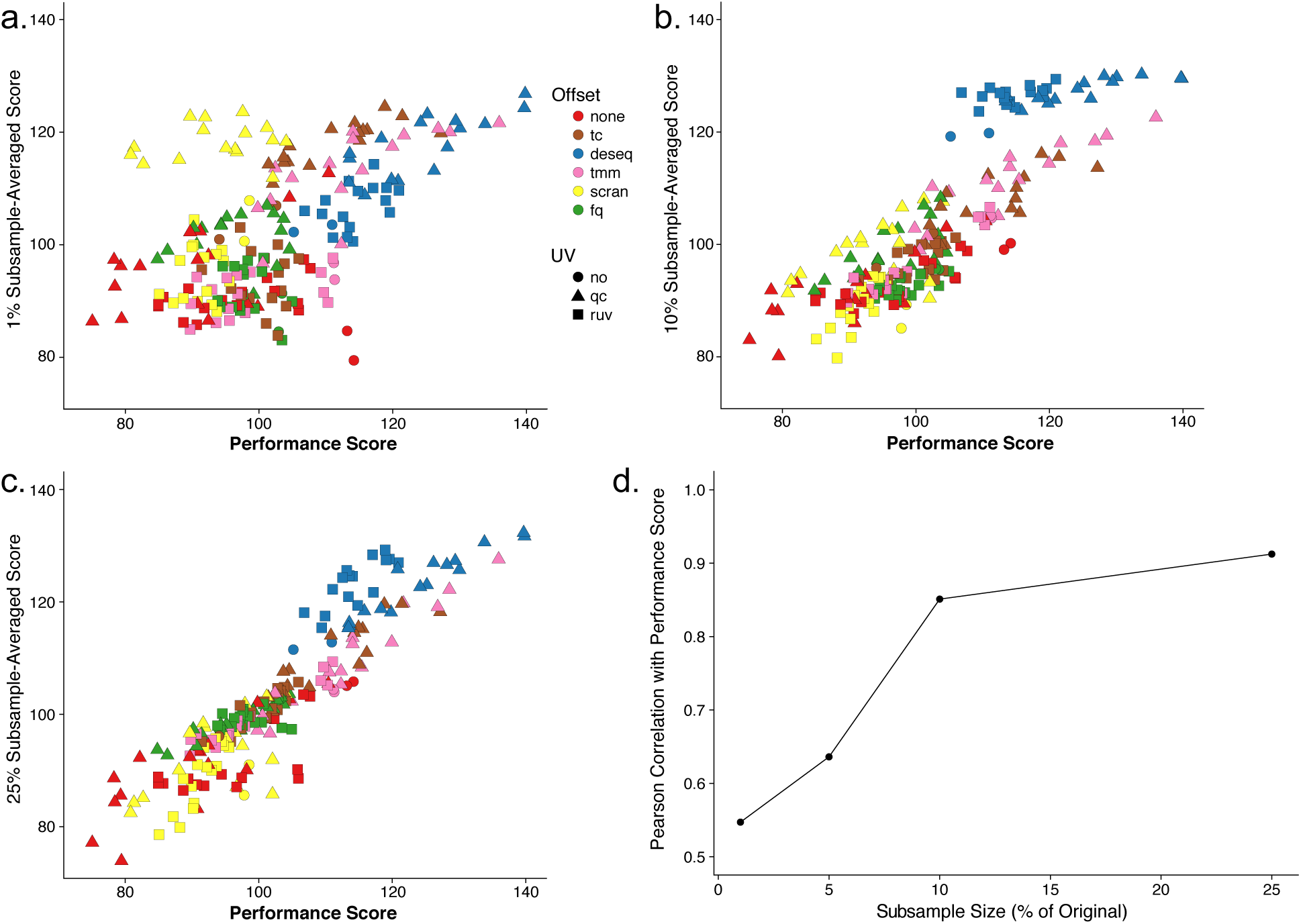
*scone* analyses for subsamples of 10x PBMC dataset [23]. (a-c) Average subsample performance score vs. full-sample performance score. We randomly extracted 10 subsamples from the full dataset corresponding to a fixed percentage of the original sample size, applied *scone* independently for each subsample, and averaged the 10 performance scores to obtain a final performance score per procedure. Plots are shown for subsamples comprising (a) 1% (b) 10%, and (c) 25% of the original sample. (d) Pearson correlation coefficient between average subsample performance score and full-sample performance score for different subsample percentages. When sampling at least 10% of the cells, we observed correlations greater than 0.8 with scores for the full data.

### External measures of differential expression validate *scone* performance

We validate *scone*’s performance assessment by relating normalized expression measures to controls derived from external differential expression studies. For the Pollen et al. dataset [22], we consider a set of positive (DE) and negative (non DE) control genes for differential expression between CP+SP (cortical plate and subplate) and SZ+VZ (subventricular zone and ventricular zone) tissues from an independent bulk microarray dataset Miller *et al*. [25]. As positive controls, we select the 1,000 most significantly differentially expressed genes from that study, as ranked by limma *p*-values [26] (all 1,000 also had *p*-value < 0.01). As negative controls, we take the 1,000 least significantly differentially expressed genes. We assess our ability to discriminate these two sets of genes in a comparison of GW16 (gestational week 16) and GW21+3 (gestational week 21, cultured for 3 weeks) cells based on the normalized Pollen *et al*. [22] dataset (using limma with voom weights [27]), generating receiver operating characteristic (ROC) curves (see Methods). Our approach identifies two clusters of procedures, including one cluster with low ROC area under the curve (AUC) and low-to-moderate *scone* performance and another cluster with high ROC AUC and moderate-to-high *scone* performance (Fig. 5a). The latter includes all FQ procedures as well as a subset of upper-quartile (UQ) procedures paired with QC adjustment. This example highlights the advantage in considering method classes rather than individual procedures, as in Figure 3d: while there is a spread in *scone* performance scores for any one scaling method, FQ performs well on average and this performance is validated by external comparisons.

**Figure 5:**
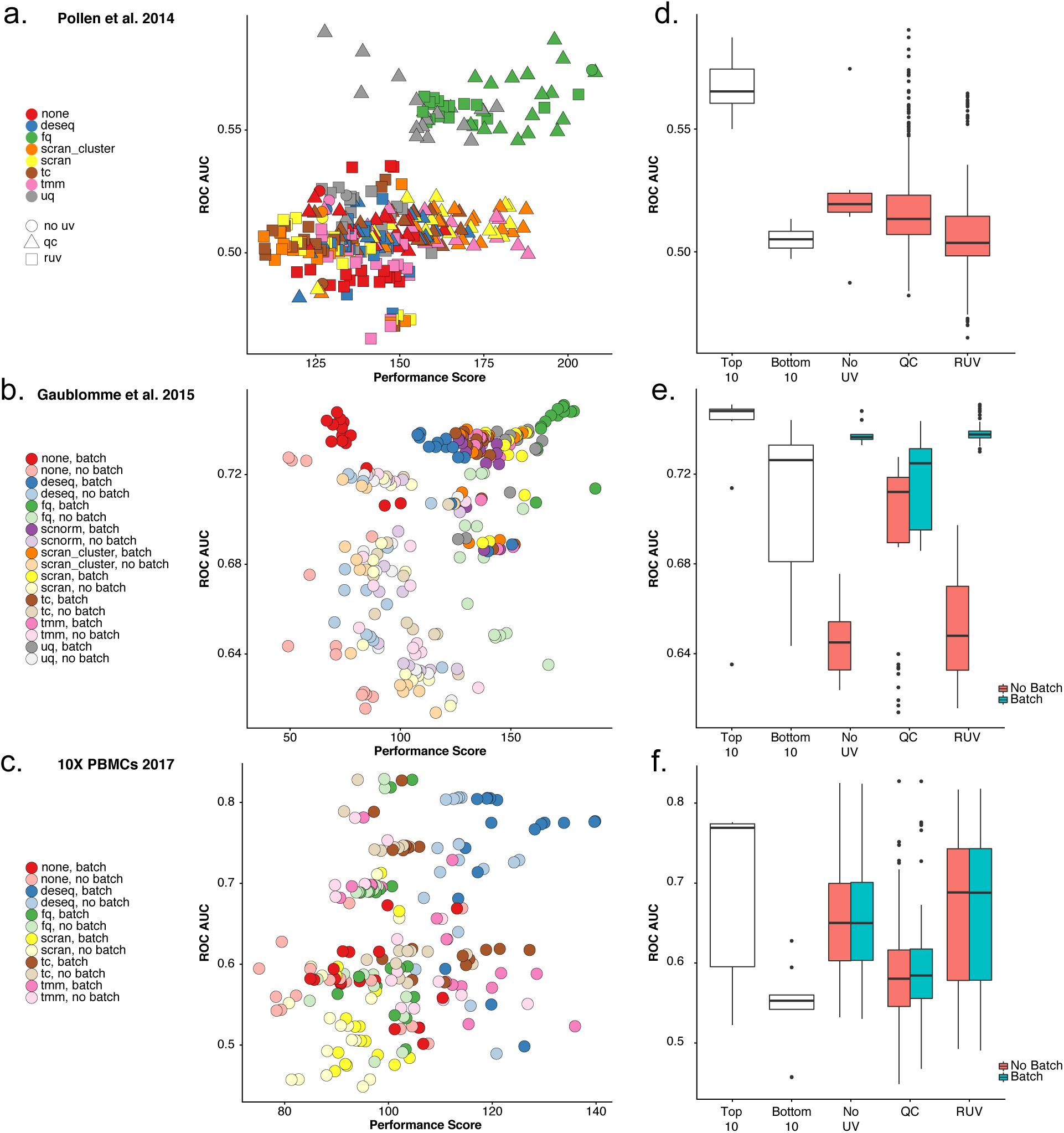
Relationship between *scone* performance scores and external differential expression validation in three scRNA-seq datasets [20, 22, 23]. (a-c) ROC AUC vs. *scone* performance score. Normalization procedures in the top-right corner are deemed best both by *scone* and by independent differential expression (DE) validation. (a) Comparing GW16 (gestational week 16) and GW21+3 (gestational week 21, cultured for 3 weeks) cells in [22], highlighting performance differences between scaling methods and the type of regression-based adjustment. (b) Comparing pathogenic and non-pathogenic cells in [20], performance differs between scaling methods and regression-based batch adjustment. (c) Comparing B cells and dendritic cells in 10x dataset [23]; performance differs between scaling methods but not by batch adjustment. (d-f) Boxplots of ROC AUC for the bottom 10 (bot10) and top 10 (top10) procedures as ranked by *scone* and for procedures with RUV, QC adjustment, and neither (“RUV”,“QC”, and “No_UV” respectively). Boxplots are further stratified by batch adjustment, when appropriate. Datasets are presented in the same order as in (a-c).

For the SMART-seq dataset of Gaublomme *et al*. [20], we utilize a separate study of bulk differential expression between pathogenic and non-pathogenic Th17 cells [28]. For each normalization procedure, we test for differences in expression between Th17-positive pathogenic cells and unsorted non-pathogenic cells, generating ROC curves. We observe a relatively low correlation between the ROC AUC and the *scone* performance scores (Fig. 5b; Spearman correlation coefficient of 0.4). However, the improved performance of both the FQ method (Fig. 3e) and our batch adjustment (Fig. 3h) are both validated by external differential expression data.

For the PBMC dataset [23], we process an independent bulk microarray dataset of Nakaya *et al*. [29]. We compute sets of positive and negative control genes by comparing baseline B cell and baseline dendritic cell (DC) microarray samples. For each normalization procedure, we use these sets to evaluate differential expression between the single-cell clusters of B cells and dendritic cells, as defined by Seurat's clustering procedure [30] (see Methods). DESeq scaling performs well on average, as suggested by the *scone* performance score (Fig. 3f).

While the *scone* ranking is not necessarily correlated with the AUC-based ranking across the whole performance range, we find an overall high level of agreement between the two rankings at the level of method classes, with the exception of RUV and QC methods. When considering many normalization procedures, users may also rely on top-ranking procedures to provide a basis for further exploration and downstream analysis. We found that the top ten normalizations as ranked by *scone* consistently performed well in terms of ROC AUC and better than procedures that consisted of scaling only (Fig. 5d-f). Taken together, these results indicate that the *scone* performance ranking is a good way of identifying suitable normalization procedures for a given dataset.

### *scone’s* normalization performance ranking is associated with improved representation of cell-cell similarity

Our validation of the *scone* performance scoring in the previous section assumes that there were different cell populations to compare. In many scRNA-seq studies, however, the goal is to identify novel cell sub-populations via cluster analysis. Here, we aim to assess the ability of *scone* to identify normalization procedures or classes thereof that will lead to the best clustering of a given dataset, using some notion of ground truth for cell clusters.

We simulate 10 datasets mimicking typical characteristics of a dataset comprising multiple cell populations using the Bioconductor R package splatter, with simulation parameters estimated from our 10x Genomics dataset (Fig. 6a; see Methods). The *scone* performance score is highly correlated with the adjusted Rand index (ARI) calculated between the simulated clusters and the clusters identified by *k*-means on the normalized data (Fig. 6b-c; see Methods).

**Figure 6:**
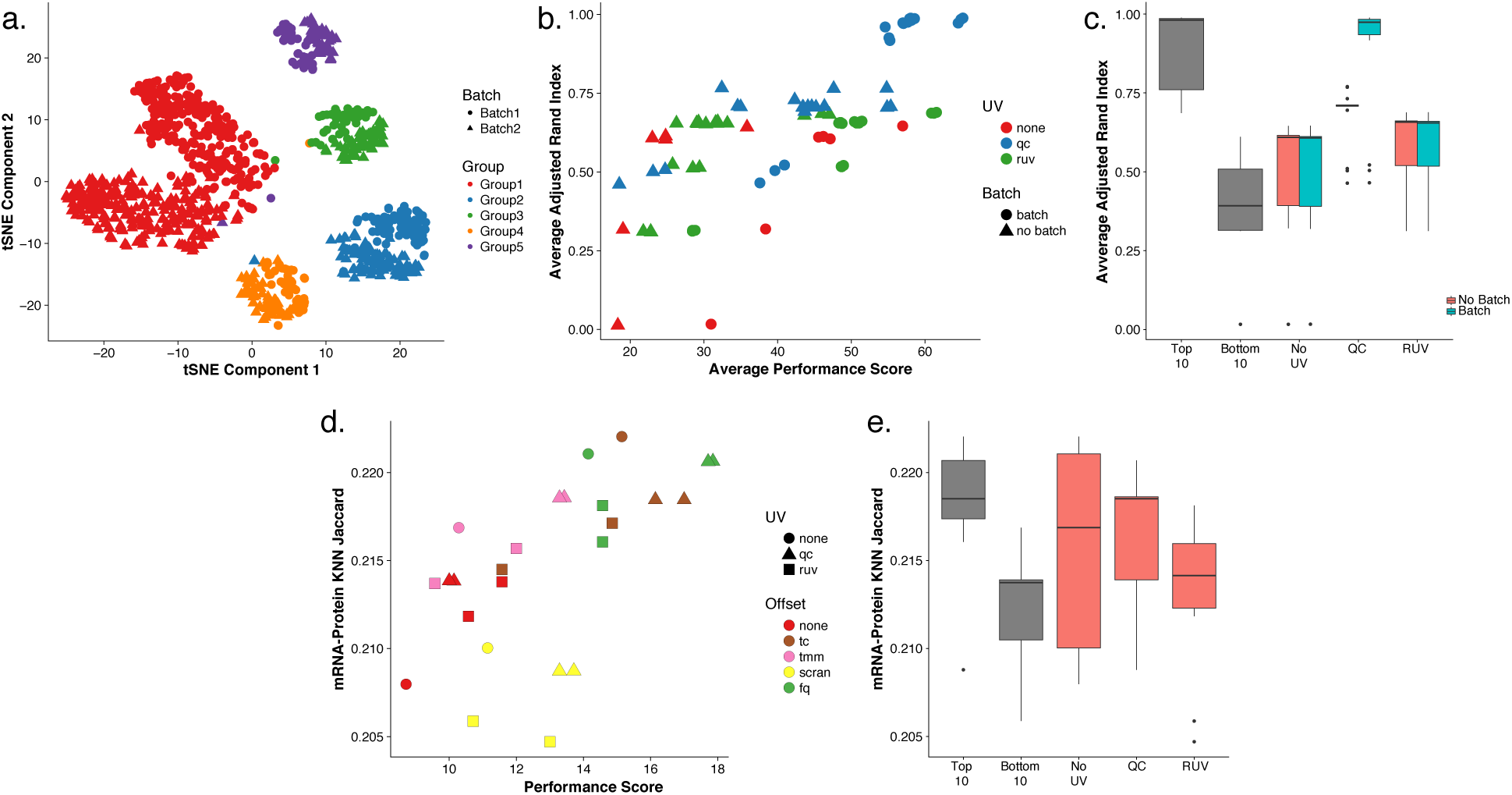
Validating *scone* performance with simulated data and external cell-level data. (a) t-distributed stochastic neighbor embedding (tSNE) of the first 10 PCs of the log-transformed, TC-normalized UMI counts for a dataset simulated using splatter, with parameters inferred from the 10x PBMC dataset [23]. (b) Average adjusted Rand index (ARI) between the true simulated clusters and *k*-means clusters (*k* = 5) for normalized data vs. *scone* performance score (without BIO_SIL score), across 10 splatter simulations (see Methods). A Pearson correlation of 0.73 between the two metrics highlights the ability of *scone* to select procedures that optimize aspects of clustering that are not explicitly accounted for in the performance panel. The top-performing procedure was FQ with adjustment for batch and 1 qPC. (c) Boxplot of average adjusted Rand index for the bottom 10 (bot10) and top 10 (top10) procedures as ranked by *scone* and for procedures with RUV, QC adjustment, and neither (“RUV”,“QC”, and “No_UV” respectively). The boxplot is stratified by batch adjustment for the latter 3 categories. (d) Jaccard score between kNN graph of protein abundance measures and kNN graph of normalized expression measures (*k* = 792, 10% of samples; see Methods) vs. *scone* performance score. A Pearson correlation of 0.60 between these metrics demonstrates how *scone* selects procedures that improve local representations of cell-cell similarity. (e) Boxplot of Jaccard score for the bottom 10 (bot10) and top 10 (top10) procedures as ranked by *scone*, procedures with no non-batch unwanted variation normalization (“No_UV”), and procedures with RUV or QC adjustment (“QC” or “RUV”).

We also apply *scone* to the recent cellular indexing of transcriptome and epitopes by sequencing (CITE-seq) dataset of Stoeckius *et al*. [31], in which gene expression and antibody levels for 13 cell-surface proteins had been jointly measured for the same cells. Specifically, we use *scone* to normalize the transcription measures and then examined the extent to which their consistency with the protein measures varies according to normalization. Computing *k*-nearest-neighbor graphs (*k =* 792, 10% of cells) for the two spaces, namely, protein abundance and transcript abundance (10 PCs, see Methods), we observe an increase in overlap between the two graphs as the *scone* performance score increased (Fig. 6d-e), reflecting how procedures ranked highly by *scone* are better at representing surface marker expression similarity.

### Using contrasts to adjust for batch effects in nested designs

The *scone* analysis of the Th17 dataset (Fig. 3) demonstrates the importance of correcting for batch effects in single-cell RNA-seq data. However, depending on the experimental design, simply including a batch variable in the model is not always a viable option. As an extreme case, imagine a completely *confounded design*, in which each biological condition is assayed in a distinct batch. In such a case, regressing out the batch indicator from the expression measures will result in the removal of biological effects; conversely, not accounting for batch will make it impossible to attribute the observed differences in expression measures to biological differences between conditions or technical differences between batches. Note that this is not just a thought experiment; several examples of datasets with suboptimal designs are discussed in Hicks *et al*. [5].

On the opposite end of the spectrum are experiments designed in such a way that each batch contains cells from each biological condition. Such *factorial designs* are the optimal choice, when possible. Hicks *et al*. [5] and Tung *et al*. [32] discuss practical aspects of designing factorial experiments in the context of scRNA-seq.

Although optimal, factorial experiments are not always possible or practical. An alternative strategy is to collect multiple batches of cells from each biological condition of interest, in what we refer to as a *nested design*. A good example of nested design is given by the iPSC dataset analyzed in Tung *et al*. [32] (Fig. 7). After scaling normalization, the cells clearly cluster by individual, but the cells for each individual are further clustered by batch (Fig. 7a). Blindly removing these batch effects with a standard batch correction method, such as ComBat [33], removes the biological effects of interest along with the batch effects (Supplementary Fig. 5). Moreover, the QC measures collected as part of the *scone* pipeline are not able to completely capture the batch effects (Fig. 7b), as the space of the first two PCs of the QC measures is dominated by the difference between a subset of low-quality cells and the rest of the cells. Explicitly accounting for the nested nature of the design while adjusting for batch effects is the only strategy that effectively removes the unwanted technical variation and preserves the biological signal of interest (Fig. 7c; see Methods for details on our nested batch effect correction).

**Figure 7:**
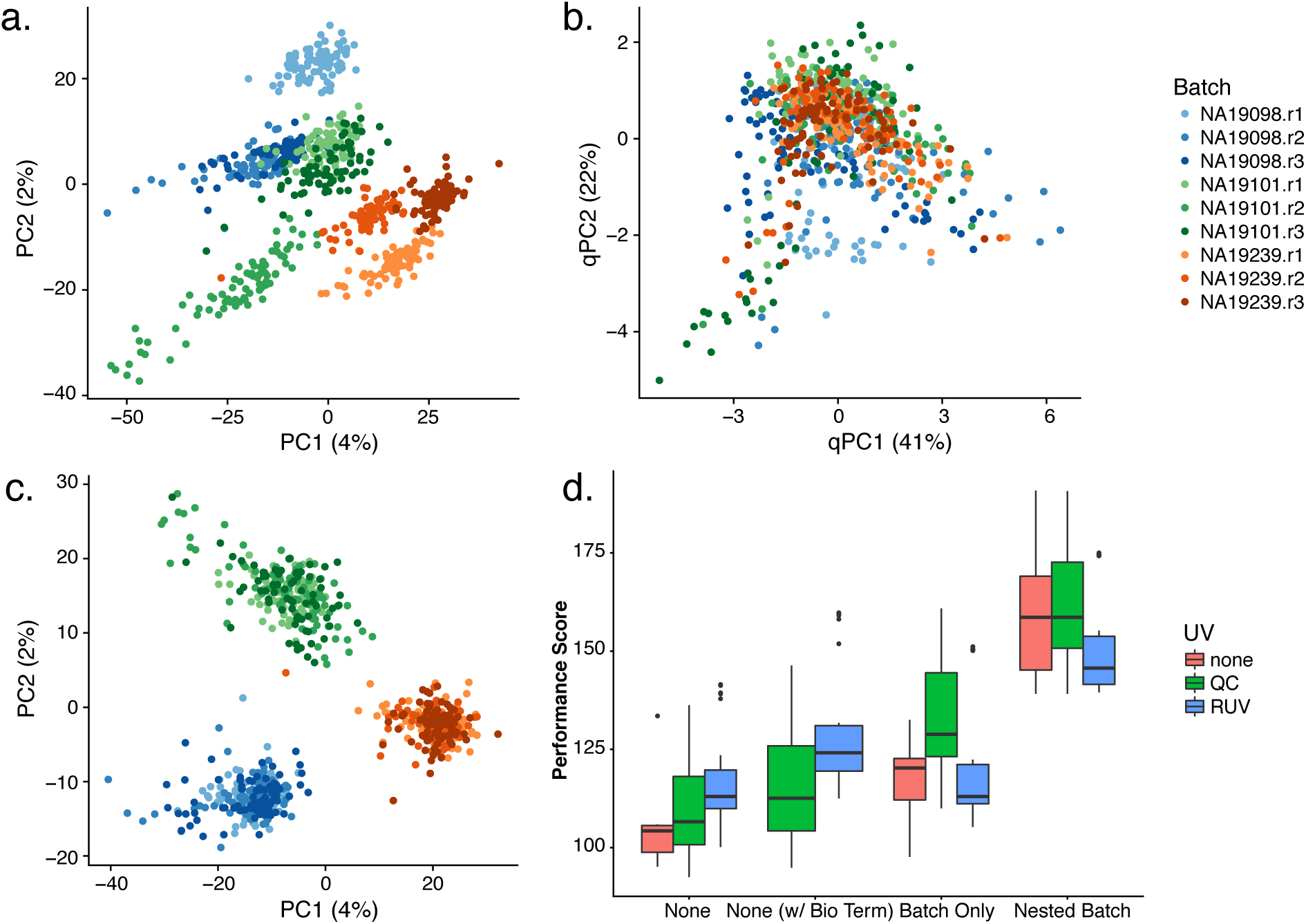
*scone* results for human induced pluripotent stem cells (iPSC) dataset with nested study design [32]. (a) PCA of the log-transformed, TC-normalized UMI counts for all genes and cells passing quality filtering, with points coded by donor (color) and batch (shade). The cells cluster by batch, indicating substantial batch effects. (b) PCA of QC measures, with points coded by donor and batch. The QC measures do not appear to capture batch effects, but rather intra-batch technical variation. (c) PCA of log-transformed expression measures after FQ normalization followed by normalization for nested batch effects (top-performing procedure in *scone*), with points coded by donor and batch. As desired, cells cluster by donor, but not by batch. (d) Boxplot of *scone* performance score, stratified by regression-based normalization. Normalization procedures including a nested batch correction performed better than those without that step.

The scone package is able to detect nested designs by examining the cross-tabulation of the biological and batch factors. Given a nested design, the nested batch effect adjustment based on Equations (3) and (4) is automatically applied as one of the different normalization strategies to be compared (see Methods). Nested designs are common in single-cell studies due to various practical constraints (e.g., processing material from different tissues separately). The *scone* performance scores (Fig. 7d) show how only procedures that remove the nested batch effects rank high in the evaluation step. This holds even when removing the two performance metrics that directly involve the batch indicator (BIO_SIL and BATCH_SIL; see Methods), suggesting that the result is not driven by these two metrics alone (Supplementary Fig. 5).

### The scone package’s user interface facilitates the exploration of normalized data

As part of the Bioconductor R package scone, we have developed a Shiny app [34] that allows users to interactively explore the data at various stages of the *scone* workflow. In Figure 8, we use the cortical neurons dataset of Pollen *et al*. [22] to illustrate the app’s functionality.

**Figure 8:**
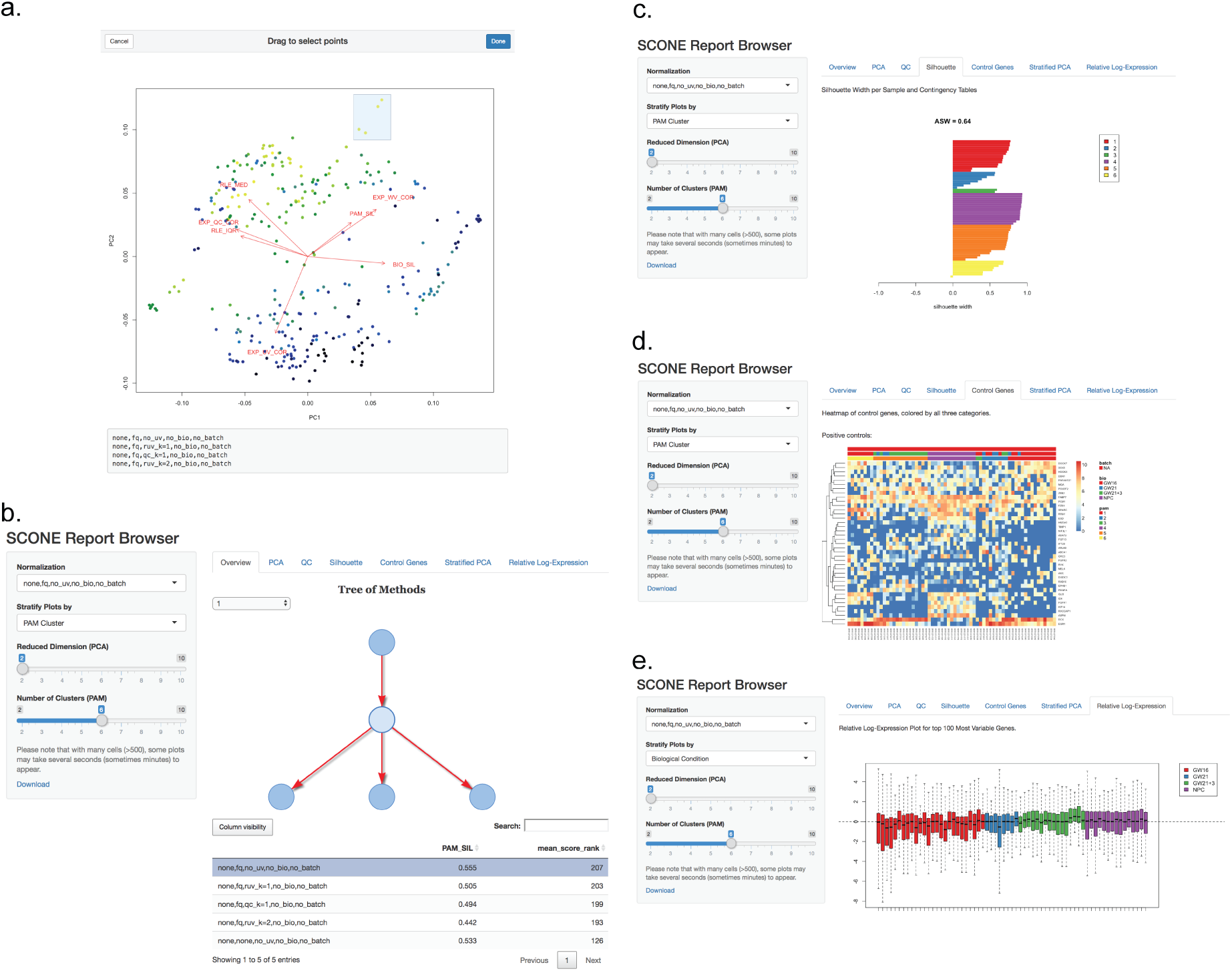
Report Browser Shiny interface. (a) Selecting normalization procedures of interest using the interactive biplot function biplot_interactive and its drag-and-drop window selection tool. This tool is useful for exploring performance clusters and selecting procedures that perform similarly across the eight performance metrics. (b) Browsing normalized products. The scone Report Browser presents an interactive tree representation (top-right panel) of selected procedures. Procedures may be further selected via a sortable performance table (bottom-right panel) or a drop-down menu (side panel). The report will then produce plots corresponding to various analyses of the normalized data. (c) Report Browser “Silhouette” tab: For the selected procedure, the silhouette width of each normalized sample is computed, grouping samples by biological condition, batch, or PAM clustering. The drop-down menu in the left bar allows the user to switch between the three categorical labels; the slider in the left panel allows the user to select the number of clusters for PAM, recomputed for each normalization procedure. (d) Report Browser “Control Gene” tab: If the user provides positive and negative control genes, the gene-level expression measures for these genes are visualized using silhouette-sorted heatmaps, including annotations for biological condition, batch, and PAM clustering. (e) Report Browser “Relative Log-Expression” tab: A boxplot of relative log-expression (RLE) measures is shown for the selected normalization procedure. Boxes (per-cell) are color-coded by biological condition, batch, or PAM clustering (drop-down selection in the left panel). If the majority of genes are not expected to be differentially expressed, the RLE distributions of the samples should be similar and centered around zero.

The scone package provides a function to display an interactive version of the biplot, allowing the user to select a group of normalizations for further exploration (Fig. 8a).

The app also provides a hierarchical overview of all the compared normalization strategies (Fig. 8b). The hierarchy is based on the series of algorithmic choices that constitute a given normalization strategy: In the example, the first level of the hierarchy represents scaling (e.g., FQ, TMM), while the second represents regression-based methods (QC or RUVg). In general, additional levels are present for the optional imputation and batch correction steps. Alternatively, the user can select a normalization strategy using the drop-down menu in the left panel of the app or using the interactive table at the bottom of the screen; this table can be sorted by the *scone* performance score or by any individual performance metric, making it easy to select, for instance, the procedure that maximizes the preservation of wanted variation (as measured by the EXP_WV_COR metric; see Methods).

Once a normalization approach has been selected for inspection, the Shiny app provides six exploratory tabs for an extended view of the normalized data, that should guide the selection of the final procedure. Here, we focus on three examples (see Methods for details). The “Silhouette” tab (Fig. 8c) shows the silhouette width for each sample, for clustering based on partitioning around medoids (PAM), clustering by batch, or clustering by biological condition (if available). If the user provides a set of positive and negative control genes, these are visualized in the “Control Genes” panel (Fig. 8d) in a heatmap that includes batch, biology, and PAM clustering information. Similarly, the “Relative Log-Expression” tab displays boxplots of the relative log-expression (RLE) measures for the normalized data (Fig. 8e).

## Discussion

Many different normalization schemes are available, either specifically designed for scRNA-seq or borrowed from the bulk RNA-seq and microarray literatures. Here, we have demonstrated that simple global-scaling normalization is not always sufficient to correctly normalize the data and that more sophisticated strategies may be needed. However, different normalization strategies may perform differently across datasets, depending on the experimental design, protocol, and platform.

The main idea behind *scone* is to use a data-driven approach to select an appropriate normalization strategy for the data at hand. Although it may be infeasible to select a “best” normalization, as this would depend on a somewhat subjective definition of optimality, *scone* provides a set of performance metrics (and clustering via the biplot) that can be used to reduce the number of normalization procedures to further explore in the selection of a suitable strategy. One advantage of a panel-based normalization selection framework is that it can be communicated and reproduced by other investigators. We have shown using real data spanning different labs and technologies that *scone* is able to reliably rank normalizations by summarizing multiple dimensions of data quality.

Given the small amount of RNA present in a single cell, scRNA-seq data are characterized by a large fraction of dropouts, i.e., genes that are expressed but not detected. To account for the resulting false negatives, *scone* estimates false negative rates that can be used to filter out low-quality samples and, optionally, to impute the expression measure of dropout genes. Zero imputation is still in its infancy; only a handful of methods exist [35, 36] and it is unclear whether their promised advantages outweigh their limitations. Although we focused on normalization, *scone* can be used to compare different filtering and imputation methods (including none). An alternative approach to imputation is to model the zeros as part of dimensionality reduction. An example of such a method, ZINB-WaVE [37], has the ability to include additional covariates to produce a low-dimensional representation of the data that is not influenced by unwanted variation. In principle, the covariates (e.g., QC or RUV factors) selected by *scone* as important for normalization can be included in ZINB-WaVE to provide a more robust projection of the data.

More generally, this manuscript discusses normalization as a modular preprocessing step, transforming read counts into normalized expression measures. Normalization can alternatively be discussed as a parameterization in model-based approaches that combine normalization with other downstream analyses, such as dimensionality reduction and differential expression [3, 19, 37]. Performance analyses of normalization as a preprocessing step can also aid such methods, for instance, by informing them on which covariates to include to adjust for unwanted technical effects.

We wish to emphasize that although *scone* has demonstrated its usefulness in published studies (e.g., Fletcher *et al*. [38], Afik *et al*. [39], Gadye *et al*. [40], and Martin-Gayo *et al*. [41]) and normalization can help remove unwanted variation from the data, a careful experimental design is the most important aspect of a successful use of scRNA-seq. In fact, if the biological effects of interest are completely confounded with unwanted technical effects, no statistical method will be able to extract meaningful signal from the data [5].

Our discussion here surrounds normalization, but the *scone* framework is more general, facilitating the comparison of imputation methods [35, 36], dimensionality reduction techniques [37, 42, 43], and additional preprocessing steps such as gene and sample filtering.

The present article also focuses on scRNA-seq, but the methodology and software are general and applicable to other types of assays, including microarray, bulk RNA-seq, adductomic, and metabolomic assays. In particular, the user could extend the package by adding different metrics for scRNA-seq, as well as metrics specific to other assays. The scone package implementation leverages core Bioconductor packages for efficient parallel computation and on-disk data representation, both essential when analyzing large datasets [44–46].

The computational complexity of *scone* is directly related to the complexity of the normalization methods included in the comparisons. In particular, all scaling methods, RUV, and regression-based methods are very efficient, leading to reasonable computing time (e.g., 60 hours with 1 CPU for the 10x Genomics PBMC dataset of 12,039 cells). With parallelization, this computation can be sped up considerably (e.g., 11 hours with 10 processors). For very large datasets, subsampling can be used to decrease computation: One or more random subsets of the samples can be used to evaluate a set of normalization procedures using the *scone* metrics and only the selected normalization can be subsequently applied to the full dataset.

Overall, *scone* provides a flexible and modular framework for the preprocessing of scRNA-seq data that can be used by practitioners to evaluate the impact of the statistical design of a given study and select an appropriate normalization, as well as by method developers to systematically compare a proposed strategy to state-of-the art approaches.

## Methods

The *scone* workflow (Fig. 2) consists of five steps: (i) Quality control; (ii) Filtering; (iii) Normalization procedures; (iv) Normalization performance assessment; (v) Exploratory analysis of normalized data. We next provide specific details for these steps. By “log” transformation, we generally refer to the ln(*x* + 1) function, so that zero counts are assigned a value of zero (R function log1p).

### Quality control

For assessing scRNA-seq data quality at the sample level, we rely on over a dozen quality control (QC) metrics evaluated by software packages such as FastQC [47], Picard [48], and Cell Ranger [49]. These QC measures summarize various aspects of genome alignment, nucleotide composition, and, for the 10x Genomics platform, unique molecular identifiers (UMI) and barcode processing. Summaries of these QC measures are presented in Tables 1 and 2.

QC measures can vary substantially between batches (e.g., C1 run) and also flag low-quality outlying samples (e.g., libraries with low percentages of mapped reads). Including these “bad” samples in downstream analyses may introduce bias in results (e.g., clustering driven by library quality). Even after sample filtering, QC measures can be strongly associated with read counts (Fig. 1). It is therefore advisable to both filter samples based on QC measures as well as consider how these measures can be used in the normalization process.

In some cases, it is difficult to obtain low-level (e.g., alignment-based) QC measures. This may occur if sequence data are filtered or omitted when uploading datasets to a public repository. A similar issue arises when expression measures are derived from simulations of count data, rather than sequence-level simulations. In either case, we may consider QC measures derived from count matrices such as those computed by the scater package [50]; we have taken this approach for our simulated data (see Datasets).

### Filtering

The goal of data filtering is to remove problematic or noisy observations from downstream analysis. This can simplify many aspects of such analyses, including normalization. In our study, we use gene and sample filtering to combat zero inflation and poor data quality. Our filtering step has three sub-steps, the latter two reducing the size of the dataset:

1. Define *common genes* based on read counts: Genes with more than *n_r_* reads in at least *f_s_* of samples, where *n_r_* is the upper-quantile of the non-zero elements of the samples-by-genes count matrix and *f_s_* is a user-specified percentage, with default value 25%.
2. Filter samples based on QC measures: Remove samples with low numbers of reads, low proportions of mapped reads (not applied to the 10x Genomics dataset), low numbers of detected common genes, or high areas under the *false negative rate* (FNR) curve as defined below. The threshold for each measure is defined data-adaptively: A sample may fail any criterion if the associated metric under-performs by *z_cut_* standard deviations from the mean metric value or by *z_cut_* median absolute deviations from the median metric value. For all sample-filtered datasets, we have used *z_cut_* = 2. This sub-step was not applied to the Pollen *et al*. [22] dataset due to the small number of samples.
3. Filter genes based on read counts: Remove genes with fewer than *n_r_* reads in at least *n_s_* samples, where *nr* is the upper-quantile of the non-zero elements of the samples-by-genes count matrix, for samples that passed the filtering described in the previous step. We have set a default of *n_s_* = 5 to accommodate markers of rare populations. This sub-step ensures that included genes are detected in a sufficient number of samples after sample filtering. Due to the small number of negative control genes in the dataset from Gaublomme *et al*. [20] (see below), we force inclusion of all detected (i.e., non-zero count) negative controls.

### False negative rate analysis

Single-cell RNA-seq data have many more genes with zero read counts than bulk RNA-seq data. In some cases, the number of zeros far exceeds predictions from standard low-mean count distributions; this zero inflation (ZI) can occur for biological reasons (i.e., the gene is simply not expressed) or technical reasons (e.g., low capture efficiency and amplification).

It is informative to examine the distribution of zero counts, as this might reveal problematic cells or genes. For instance, cells with an unusually high proportion of zero counts for highly-expressed genes might have encountered PCR or read alignment problems. Additionally, in an attempt to determine whether zeros are technical, one can employ *housekeeping genes* which are expected to be expressed both highly and uniformly across cells.

Specifically, let *Y_ij_* denote the log-read count of gene *j* ∈ {1,…, *J*} in cell/sample *i* ∈ *{1,…, n}* and let 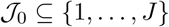 denote a set of housekeeping genes. For each cell *i*, fit the following *logistic regression* model to the housekeeping genes only:

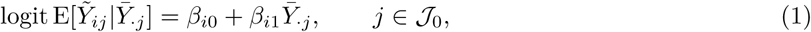

where 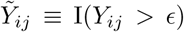 is an indicator variable equal to 1 if gene *j* is “detected” and zero otherwise (as determined by a detection threshold ∊ > 0), 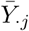 is the median log-read count of gene *j* across all nonzero samples, *β_i_* = (*β_i_*_0_*,β_i_*_1_) ∈ ℝ^2^ are cell-specific regression parameters, and the logit function is defined as 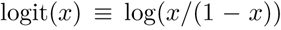 for *x* ∈ [0,1]. The motivation behind this model is that the detection rate 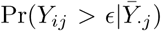 of gene *j* in cell *i* should be related to the “baseline” expression measure 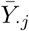 of the gene in a cell-specific manner (i.e., cell-specific regression parameters *β*_i_) and that undetected housekeeping genes should constitute “false negatives”. In practice, we set ∊ = 0.

For a given detection threshold *∊*, Equation (1) yields FNR curves for each cell, where each point on the curve corresponds to a gene’s “failure probability” 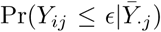 at a baseline expression 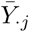. The higher the curve, the lower the quality of the sample. Samples can then be compared based on the *area under the curve* (AUC) for their respective FNR curves.

Note that we assume that the housekeeping genes are both highly and uniformly expressed across all samples. These assumptions allow us to treat the zeros as “dropouts” rather than as biologically meaningful zeros, hence the interpretation of 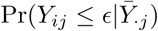 as failure probability.

## Normalization procedures

A variety of gene- and sample-specific unwanted effects can bias gene expression measures and thus require normalization. Accordingly, we distinguish between two types of normalization: Within-sample normalization [51], which adjusts for gene-specific (and possibly sample-specific) effects, e.g., related to gene length and GC-content, and between-sample normalization, which adjusts for effects related to distributional differences in read counts between samples, e.g., sequencing depth, C1 run, library preparation. Here, we focus on between-sample normalization.

Most between-sample normalization methods proposed to date are adaptations of methods for bulk RNA-seq and microarrays and range, as described next, from simple global scaling to regression on gene- and sample-level covariates.

### Global-scaling normalization

For the simplest linear *global-scaling normalization* procedures, gene-level read counts are scaled by a single factor per sample.

**Total-count (TC)**. The scaling factor is the sum of the read counts across all genes, as in the widely-used reads per million (RPM), counts per million (CPM), and reads per kilobase of exon model per million mapped reads (RPKM) [7].

**Single-quantile**. The scaling factor is a quantile of the gene-level count distribution, e.g., upper-quartile (UQ) [52].

**Trimmed mean of M values (TMM) [8]**. The scaling factor is based on a robust estimate of the overall expression fold-change between the sample and a reference sample. TMM is implemented in the Bioconductor R package edgeR [53]. The default behavior used here is that the selected reference sample has an upper quartile closest to the mean upper quartile of all samples.

**DESeq [9]**. The scaling factor for a given sample is defined as the median fold-change between that sample and a synthetic reference sample whose counts are defined as the geometric means of the counts across samples. The method is implemented in the Bioconductor R packages DESeq and edgeR (as “RLE”) [53, 54]. Note that the method discards any gene having zero count in at least one sample; as zeros are common in single-cell data, the scaling factors are often based on only a handful of genes.

**scran [14]**. To reduce the effect of single-cell noise on normalization, the scaling factors are computed on pooled expression measures and then deconvolved to obtain cell-specific factors. The method is implemented in the Bioconductor R package scran [55]. Pool sizes from 20 to the 100 (intervals of five) were considered. For the dataset of Pollen *et al*. [22], we limit pool size to the total number of cells.

Optionally, cells can be clustered prior to normalization to relax the assumption that the majority of genes are not differentially expressed across groups of cells (“scran_cluster” scaling). In these cases, we used the quick clustering utility in scran, setting a minimum cluster size at 50. Minimum cluster size was then used as the pool size.

While not strictly necessary, our implementation of global-scaling procedures typically preserves the original measurement scale by further rescaling the normalized measures by the inverse mean scaling factor.

### Non-linear scaling normalization

In some cases, a single scaling factor per sample may not be sufficient to capture the non-linear effects that affect gene expression measures. Hence, some authors have proposed non-linear normalization methods. Although, these are not technically “scaling” methods, they belong to the first step of the *scone* normalization template, in that they are aimed at making the between-sample distributions of expression measures more similar, rather than explicitly correcting for batch or other confounding factors.

**Full-quantile normalization (FQ)**. All quantiles of the read count distributions are matched between samples [52]. Specifically, for each sample, the distribution of sorted read counts is matched to a reference distribution defined in terms of a function of the sorted counts (e.g., median) across samples. This approach, inspired from the microarray literature [56], is implemented in the Bioconductor R package EDASeq [57].

**SCnorm [17]**. Bacher *et al*. [17] noted that a single scaling factor per sample is not enough to account for the systematic variation in the relationship between gene-specific expression measures and sequencing depth. To address this problem, they use quantile regression to estimate the dependence of gene expression measures on sequencing depth, group genes with similar dependence, and use a second quantile regression to estimate scaling factors within each group. In this way, gene expression measures are normalized differently across the range of expression (i.e., highly-expressed genes are scaled differently than lowly-expressed genes). The method is implemented in the Bioconductor R package SCnorm [58]).

### Regression-based normalization

We consider the following *generalized linear model* (GLM; Figure 2), which allows adjustment for known and unknown factors of “unwanted variation”:

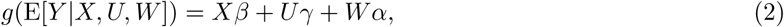

where *Y* is the *n* × *J* matrix of gene-level read counts, *X* is an *n* × *M* design matrix corresponding to the *M* covariates of interest/factors of “wanted variation” (e.g., treatment) and *β* its associated *M* × *J* matrix of parameters of interest, *U* is an *n* × *H* matrix corresponding to known factors of unwanted variation (e.g., batch, sample QC measures) and *γ* its associated *H* × *J* matrix of nuisance parameters, *W* is an *n* × *K* matrix corresponding to unknown factors of unwanted variation and *α* its associated *K* × *J* matrix of nuisance parameters, and *g* is a link function, such as the logarithm in Poisson/log-linear regression.

The *Uγ* and *Wα* terms correspond, respectively, to supervised and unsupervised removal of unwanted variation. A fully supervised version of Equation (2), without *W*, reduces to a simple GLM fit. The remove unwanted variation (RUV) model of Risso *et al*. [11] arises as a special case of Equation (2), when one omits the known unwanted factors *U*. In many cases, the data-driven unsupervised version of Equation (2), without *U*, captures effects related to *U*; for instance, *W* is often associated with QC measures (Supplementary Fig. 4). However, in many cases, *U* should still include known batches (e.g., set of samples processed at the same time), as *W* could capture effects related to sample quality within batches that are important to remove in addition to the batch effects captured by *U*. We also find that, in practice, the computationally simpler approach of fitting a linear model to log-counts *Y* yields good results. Hence, *scone* relies on a linear model version of Equation (2) with identity link function (*g*(*x*) = *x*).

As detailed in Risso *et al*. [11] and implemented in the Bioconductor R package RUVSeq [59], the unknown, unwanted factors *W* can be estimated by singular value decomposition (SVD) using three main approaches. Here, we only use RUVg, which estimates the factors of unwanted variation based on *negative control genes*, assumed to have constant expression across all samples (*β* = 0).

In the current implementation of scone, the only covariates that can be included in *U* are those related to known experimental batches and to sample-level QC. Because QC measures are often highly correlated (Fig. 1), scone transforms the metrics using principal component analysis prior to fitting the model. The user can choose the maximum number of components to include in the model.

Single-cell RNA-seq data can exhibit severe zero inflation (especially in the case of the 10x Genomics platform, see Supplementary Fig. 1c): anywhere from 10–90% of the read counts are zero (Supplementary Fig. 1i). Regression-based normalization may modify these zeros, rendering their interpretation problematic (cf. technical or biological contributions to zero inflation). To avoid these issues, we restore all values that were initially zero back to zero following normalization. We also make sure that any negative values produced by a normalization procedure are set to zero. As a result, the performance metrics do not reflect differences in the effects of normalization on zero values.

### Adjustment for nested experimental designs

The model of Equation (2) should be applied with caution and only after a careful examination of the experimental design. In particular, a common limitation of scRNA-seq datasets is the *nesting* of unwanted technical effects within the biological effects of interest. For instance, the iPSC dataset of Tung *et al*. [32] contains samples derived from three individuals, each processed in three batches. Regressing read counts on the batch covariate in *U* without adjusting for the covariate of interest in *X* would then remove the effect of interest (effect of individual donor). Additionally, to avoid collinearity issues between columns of *U* and *X* due to nesting, one could either specify suitable contrasts or use a mixed effect model where technical effects are viewed as random. The scone implementation of the model in Equation (2) automatically detects nested designs and specifies the right contrasts to avoid removing the effects of interest and to avoid identifiability issues in the estimation procedure. See Tung *et al*. [32] for an alternative method that uses random effects.

Specifically, to account for nested batch effects, we consider the following model for each gene (cf. ANOVA). For illustration purposes and ease of notation, we do not include the gene subscript and the additional known or unknown covariates allowed in Equation (2), although the latter can be included in the scone package implementation. Let *Y_ijk_* denote the expression measure of sample *k* in batch *j* of condition *i*, with *i* = 1*,…, a* conditions, *j* = 1*,…, b_i_* batches for condition *i*, and *k* = 1, …, *n_j_*_(_*_i_*_)_ samples for batch *j* of condition *i*. We fit the following model gene by gene

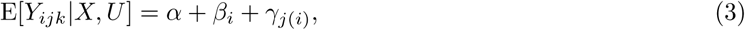

where *α* is an intercept, *β_i_* a biological effect of interest, and *γ_j_*_(_*_i_*_)_ a nested batch effect. Given the *a* + 1 constraints

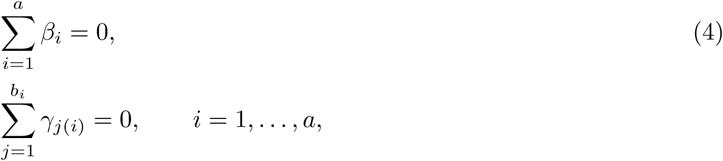

the model is identifiable and can be fit using standard R functions such as lm. The batch-corrected gene expression measures are then given by the residuals *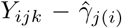*. When additional factors of unwanted variation are included in the model, their effects are similarly subtracted from the original matrix to produce the normalized matrix.

Note that although *scone* is able to handle the special case of nested designs, no adjustment method is able to remove batch effects while preserving biological effects of interest if the experimental design is completely confounded. For instance, if only one batch had been processed per individual in the iPSC dataset of Tung *et al*. [32], it would have been impossible to determine if the differences between samples were due to batch effects or biology [5].

## Normalization performance assessment

### Seurat clustering analyses

It is common to see clear biological clustering at early stages of a single-cell analysis (e.g., major blood cell types). However, an important asset of single-cell approaches is their ability to resolve deeper and more subtle biological heterogeneities. Thus, one might wish to maintain the large-scale clustering evident in loosely normalized data, by passing this clustering to scone as a biological classification to be preserved after normalization.

We took this approach for our two largest datasets, namely, the 10x PBMC and CITE-seq CBMC datasets. After sample filtering, we loaded the UMI matrices for these datasets into the widely-used Seurat analysis pipeline [30]. Following TC normalization, log-transformation, scaling, and principal component analysis (PCA), we clustered the cells in the first 10 principal components (PCs) at a “resolution” of 0.6. The resulting clusters were treated as biological conditions for evaluating the biological cluster tightness (BIO_SIL, see below).

For the PBMC dataset, we manually collapsed clusters based on expression of the following marker genes: IL7R (CD4 T-cells), CD14 and LYZ (CD14+ Monocytes), MS4A1 (B cells), CD8A (CD8 T-cells), FCGR3A and MS4A7 (FCGR3A+ Monocytes), GNLY and NKG7 (NK-cells), FCER1A and CST3 (Dendritic cells), and PPBP (Megakaryocytes). These RNA markers were discussed as part of the Seurat vignette.

### Normalization performance metrics

Different normalization procedures can lead to vastly different distributions of gene expression measures. We have previously shown, in the context of bulk RNA-seq, that the choice of normalization procedure had a greater impact on differential expression results than the choice of DE test statistic [52]. A natural and essential question is therefore whether normalization is beneficial and, if so, which method is most appropriate for a given dataset. In order to address this question, we have developed eight normalization performance metrics that relate to three aspects of the distribution of gene expression measures.

**Clustering properties**. The following three metrics evaluate normalization procedures based on how well the samples can be grouped according to factors of wanted and unwanted variation: Clustering by wanted factors is desirable, while clustering by unwanted factors is undesirable.

As clustering quality measures, we use silhouette widths [60]. For any clustering of *n* samples, the *silhouette width* of sample *i* is defined as

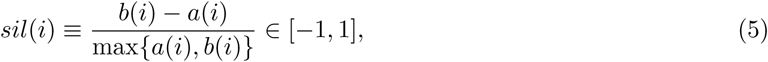

where *a*(*i*) denotes the average distance (by default, Euclidean distance over first three PCs of expression measures) between the *i*th sample and all other samples in the cluster to which *i* was assigned and *b*(*i*) denotes the minimum average distance between the *i*th sample and samples in other clusters. Intuitively, the larger the silhouette widths, the better the clustering. Thus, the *average silhouette width* across all *n* samples provides an overall quality measure for a given clustering.

- *BIO_SIL:* Group the *n* samples according to the value of a categorical covariate of interest (e.g., known cell type, genotype) and compute the average silhouette width for the resulting clustering.
- *BATCH_SIL:* Group the *n* samples according to the value of a nuisance categorical covariate (e.g., batch) and compute the average silhouette width for the resulting clustering.
- *PAM_SIL:* Cluster the *n* samples using *partitioning around medoids* (PAM) for a range of user-supplied numbers of clusters and compute the maximum average silhouette width for these clusterings. *scone* provides an option for stratified scoring, computing the PAM_SIL metric in all distinct strata defined jointly by biological and batch classification. The reported PAM_SIL metric is a weighted average across all strata, weighting by the total number of cells in each. This option is useful when prior biological classifications are poor proxies for single-cell states, i.e., when additional heterogeneity is expected. We used this option for all datasets but the 10x Genomics PBMC and CITE-seq CBMC datasets for which the biological classification were data-derived clusters (see above Seurat clustering analyses).

Large values of BIO_SIL and PAM_SIL and low values of BATCH_SIL are desirable.

**Association with control genes and QC metrics**. The next three metrics concern the association of log-count principal components (default 3) with “evaluation” principal components of wanted or unwanted variation.

- *EXP_QC_COR:* The weighted coefficient of determination 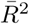 (details below) for the regression of logcount principal components on all principal components of user-supplied QC measures (using prcomp with scale=TRUE; QC measures described in Tables 1 or 2).
- *EXP_UV_COR:* The weighted coefficient of determination 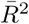 for the regression of log-count principal components on factors of unwanted variation UV (default 3) derived from negative control genes (preferably different from those used in RUV). The submatrix of log-transformed unnormalized counts for negative control genes is row-centered and scaled (i.e., for each row/gene, expression measures are transformed to have mean zero and variance one across columns/cells) and factors of unwanted variation are defined as the right-singular vectors as computed by the svds function from the rARPACK package.
- *EXP_WV_COR:* The weighted coefficient of determination 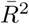 for the regression of log-count principal components on factors of wanted variation WV (default 3) derived from positive control genes. The WV factors are computed in the same way as the UV factors above, but with positive instead of negative control genes.

Large values of EXP_WV_COR and low values of EXP_QC_COR and EXP_UV_COR are desirable.

The weighted coefficients of determination are computed as follows. For each type of evaluation criterion (i.e., QC, UV, or WV), regress each expression PC on all supplied evaluation PCs. Let *SST_k_*, *SSR_k_*, and *SSE_k_* denote, respectively, the total sum of squares, the regression sum of squares, and the residual sum of squares for the regression for the *kth* expression PC. The *coefficient of determination* is defined as usual as

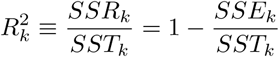

and our weighted average coefficient of determination as

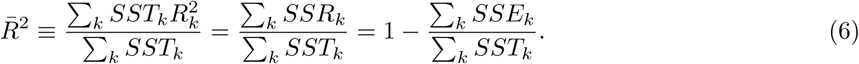

**Global distributional properties**. When comparing distributions of expression measures between samples, gene-level *relative log-expression* (RLE) measures, defined as log-ratios of read counts to median read counts across samples, are more informative than log-counts [61]:

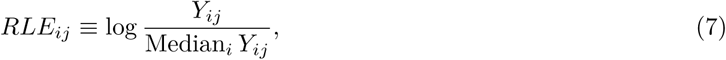

for gene *j* in cell *i*. For similar distributions, the RLE should be centered around zero and have have similar spread across samples.

- *RLE_MED:* Mean squared median relative log-expression.
- *RLE_IQR:* Variance of inter-quartile range (IQR) of RLE.

Low values of RLE_MED and RLE_IQR are desirable.

**Ranking and selecting normalization procedures**. In the *scone* framework, the expression measures are normalized according to a user-specified set of methods (including no normalization) and the eight metrics above are computed for each normalized dataset.

The performance assessment results can be visualized using biplots [24] and the normalization procedures ranked based on a function of the performance metrics. In particular, we define a *performance score* by orienting the metrics (multiplying by ±1) so that large values correspond to good performance, ranking procedures by each metric, and averaging the ranks across metrics.

Note that a careful, global interpretation of the metrics is recommended, as some metrics tend to favor certain methods over others, e.g., EXP_UV_COR naturally favors RUVg, especially when the same set of negative control genes are used for normalization and evaluation. We have used non-overlapping sets of control genes for all of the analyses discussed here.

**Performance assessment on subsampled datasets**. For the analysis presented in Figure 4, we have randomly and independently drawn 10 subsets of cells, running *scone* separately on each subset, and averaging the *scone* performance scores across the 10 subsets. We present results over a range of subsample sizes (1%, 5%, 10%, and 25% of cells) and correlate subsampled performance scores with performance scores derived from the full data.

### Evaluation of *scone* rankings

To independently evaluate the *scone* normalization performance metrics, we compared the *scone* rankings to rankings based on external data available for our three primary datasets, a CITE-seq dataset, and simulated data.

**SMART-seq C1 Th17 dataset [20]**. To evaluate *scone* performance on the Th17 dataset, we analyzed independent bulk microarray data from Lee *et al*. [28], available on GEO with accession GSE39820. We computed sets of positive (DE) and negative (non DE) control genes by comparing cytokine activation IL-1beta, IL-6, IL-23 and cytokine activation TGF-beta1, IL-6 microarray samples, using the Bioconductor R package limma [26]. Specifically, we identified as positive controls the 1,000 genes with the smallest *p*-values and as negative controls the 1,000 genes with the largest *p*-values on the microarray data, excluding any *scone* controls from these sets. For each normalization procedure, we then applied limma with voom weights [27] to the scRNA-seq data to perform differential expression analysis between Th17-positive pathogenic cells and unsorted non-pathogenic cells. We used the resulting *p*-values to infer the positive and negative controls derived from the microarray data and produce a *receiver operating characteristic* (ROC) curve. The ranking of normalization procedures by *scone* was then compared to the ranking by the ROC area under the curve (AUC).

**Fluidigm C1 cortex dataset [22]**. We similarly processed independent bulk microarray data from Miller *et al*. [25], available from the BrainSpan atlas (http://brainspan.org/static/download.html). We computed sets of positive (DE) and negative (non DE) control genes by comparing CP+SP and SZ+VZ tissues, using the limma package [26] as for the Th17 dataset above. For each normalization procedure, we then applied limma with voom weights [27] to the scRNA-seq data to perform differential expression analysis between the GW16 and GW21+3 conditions. The ranking of normalization procedures by *scone* was compared to the ranking by the ROC AUC, as detailed for the Th17 dataset.

**10x Genomics PBMC dataset [23]**. We processed independent bulk microarray data from Nakaya *et al*. [29], available on GEO with accession GSE29618. We computed sets of positive (DE) and negative (non DE) control genes by comparing baseline B cell and baseline dendritic cell (DC) microarray samples, using the limma package [26] as for the Th17 dataset above. For each normalization procedure, we then applied limma with voom weights [27] to the scRNA-seq data to perform differential expression analysis between the B cell and dendritic cell clusters, as defined by Seurat [30]. The ranking of normalization procedures by *scone* was compared to the ranking by the ROC AUC, as detailed for the Th17 dataset.

**CITE-seq dataset [31]**. Antibody-associated cellular indexing of transcriptome and epitopes by sequencing (CITE-seq) UMI counts were extracted from GEO entry GSE100866 (CBMC_8K_13AB_10X), for a collection of human cord blood mononuclear cells (CBMC) and mouse cells. Cell UMI profiles were transformed using the centered log-ratio. Means and standard deviations for each of the 13 antibody measures were computed across all mouse samples (< 0.1 human RNA UMI fraction, see Datasets) and the mean plus the standard deviation was subtracted from all abundances, as in the CITE-seq manuscript. For human samples (see Datasets), we constructed a *k*-nearest-neighbor (kNN) graph using the Euclidean metric in 13-dimensional antibody space (*k* = 792 or 10% of all human samples). For each normalization procedure, we applied PCA to log-transformed scRNA-seq data, selected the top 10 PCs, and used them to constructed a kNN graph with the same choice of *k*. The ranking of normalization procedures by *scone* was then compared to the ranking by the Jaccard similarity score of the RNA and protein kNN adjacency matrices. Note that we considered other values for *k* (e.g., *k* = 8 or 1% of samples; data not shown), but found that the mean and range of Jaccard similarity scores decreased considerably for smaller neighborhood sizes. Mouse cells were not utilized beyond preprocessing.

**Simulated datasets**. We simulated 10 independent datasets (20,000 genes by 1,000 cells each) using the Bioconductor R package splatter [62]. Simulation parameters were inferred from a subset of 100 cells from the 10x Genomics “pbmc4k” dataset [23], setting the differential expression probability to 0.3 and adding five cell populations (or “groups”) of different sizes: one population comprising 50% of the cells, one comprising 20%, and the remaining three populations comprising 10% of cells each. We switched on dropouts and added a batch effect (two batches of 500 cells) to make normalization more challenging.

We ran *scone* on each simulated UMI dataset, including a sample filtering scheme similar to the one above and additionally requiring at least 1,000 UMIs, greater than 80% of common genes detected, below 0.65 AUC, and using a *z_cut_* of 3 for greater data-adaptive leniency. Negative control genes (200 for evaluation) were perfect (log-fold change of zero) and extracted from the simulation. 200 positive control genes were selected based on maximum absolute average fold-change as reported by the simulation.

The scater package [50] was used to compute the following QC measures per simulated library: *log*10 total UMIs, *log*10 total UMI features, percent UMIs in top 50, 100, 200, and 500 features. Batch information – not group information – was extracted from the simulation. We considered procedures with zero to three factors of QC or RUVg. We used PCA to decompose the log-normalized count matrix following each normalization. We then performed *k*-means clustering on the space of the first 10 principal components, specifying the right number of clusters (*k* = 5). Finally, we computed the adjusted Rand index (ARI) between the true simulated clusters and the clusters inferred by *k*-means. We reported the average ARI across the 10 simulated datasets.

### Bioconductor R software package scone

Our *scone* framework is implemented in the open-source R package scone, freely available through the Bioconductor Project [44] at https://bioconductor.org/packages/scone.

The package implements methods for sample and gene filtering, provides wrappers for commonly-used normalization procedures, and allows the user to define custom normalization methods in the form of simple R functions.

The main function in the package, called scone, can be used to run and compare different normalization strategies. Through its integration with the BiocParallel [45] and rhdf5 [46] packages, scone can take full advantage of parallel processing and on-disk data representation, to avoid storing all the normalization results in memory and instead store the results in HDF5 files. See the package vignette at https://bioconductor.org/packages/scone for additional details.

Although here we used PCA to evaluate the performance of normalization methods (in particular, regarding the metrics EXP_QC_COR, EXP_UV_COR, and EXP_WV_COR), the scone package allows the choice of other dimensionality reduction techniques. scone also has an optional imputation step that can be turned on to evaluate the performance of imputation methods and their effect on normalization.

In addition to the ability to retrieve normalized expression measures, scone allows the user to retrieve the related design matrix. This is useful for downstream differential expression analysis, where rather than removing the unwanted variation and using the residuals as expression measures, one may want to include the factors of unwanted variation as additional covariates in a regression model.

### Datasets

**SMART-seq C1 Th17 dataset [20]**. Cells were harvested from two C57BL/6J and three IL-17A – GFP+ mice. Unsorted non-pathogenic cells were collected from the first two mice and both IL-17A-sorted pathogenic and non-pathogenic cells were collected from the three remaining mice. Cells were sorted and a Fluidigm C1-based SMART-seq protocol was used for single-cell RNA extraction and sequencing. Following sample filtering, 337 cells were retained from four donor mice – one mouse was filtered out due to the small number of acceptable cells. We consider the problem of normalizing 7,590 gene features over these 337 cells. For *scone*, we provided negative and positive control genes based on Supplementary Table S6 from Yosef *et al*. [63].

**Fluidigm C1 cortex dataset [22]**. 65 cells from the developing cortex were assayed using the Fluidigm C1 microfluidics system. Each cell was sequenced at both high and low depths; we focus on the high-coverage data. The data are available as part of the Bioconductor R package scRNAseq (https://bioconductor.org/packages/scRNAseq). No sample filtering was applied to this dataset and 4,706 genes were retained following gene filtering. For *scone*, we provided default negative control genes from the “housekeeping” list and positive control genes related to neurogenesis as annotated in MSigDB (JEPSEN_SMRT_TARGETS and GO NEURAL PRECURSOR CELL PROLIFERATION; http://software.broadinstitute.org/gsea/msigdb/cards/).

**Fluidigm C1 iPSC dataset [32]**. Three batches of 96 libraries from each of three YRI iPSC lines were sequenced using the Fluidigm C1 microfluidics system. The full dataset, including UMI counts, read counts, and quality metrics, was obtained from https://github.com/jdblischak/singleCellSeq. Library-level QC measures included: (i) Proportion of reads aligning to ERCC spike-ins (matching the pattern “ÊRCC”); (ii) number of unique molecules; (iii) well number as reported in online metadata (“wel”); (iv) concentration as reported in online metadata (“concentration”); (v) number of detected molecule classes (genes with more than zero UMI). Following gene and sample filtering, we retained 6,818 genes and 731 libraries; retained samples had more than 24,546 reads, more than 80% of common genes detected, and FNR AUC below 0.65. For *scone*, we provided default negative control genes from the “housekeeping” list, as well as positive control genes from the “cellcycle_genes” default list. ERCC genes were used as negative controls for RUVg normalization. Patient was used as a proxy for biological condition, while batch was defined as an individual C1 run.

**10x Genomics PBMC dataset [23]**. We considered scRNA-seq data from two batches of peripheral blood mononuclear cells (PBMC) from a healthy donor (4k PBMCs and 8k PBMCs). The data were downloaded from the 10x Genomics website (https://www.10xgenomics.com/single-cell/) using the cellrangerRkit R package (v. 1.1.0). After filtering, 12,039 cells and 10,310 genes were retained. For *scone*, we provided default negative control genes from the “housekeeping” list and positive control genes as the top 513 most common genes annotated in the MSigDB C7 immunological signature collection (http://software.broadinstitute.org/gsea/msigdb/collections.jsp). Seurat-derived clusters [30] were used as a biological condition, while batch was defined as an individual 10x run.

**CITE-seq dataset [31]**. One cellular indexing of transcriptome and epitopes by sequencing (CITE-seq) dataset was extracted from GEO entry GSE100866 (CBMC_8K_13AB_10X), for a collection of human cord blood mononuclear cells (CBMC) and mouse cells. 8,005 cells were called as human based on greater than 90% human-mapped UMI fraction. After filtering, 7,978 cells and 7,231 genes were retained. For *scone*, we provided default negative control genes from the “housekeeping” list and positive control genes as the top 513 most common genes annotated in the MSigDB C7 immunological signature collection (http://software.broadinstitute.org/gsea/msigdb/collections.jsp). Seurat-derived clusters [30] were used as a biological condition. QC features were limited to the fraction of human, mouse, and ERCC UMIs (3 features).

For the Th17 and cortex datasets, SRA-format files were downloaded from the Sequence Read Archive (SRA) and transformed to FASTQ format using the SRA Toolkit. Reads were aligned with TopHat (v. 2.0.11) [64] to the appropriate reference genome (GRCh38 for human samples, GRCm38 for mouse). RefSeq mouse gene annotation (GCF_000001635.23_GRCm38.p3) was downloaded from NCBI on Dec. 28, 2014. RefSeq human gene annotation (GCF_000001405.28) was downloaded from NCBI on Jun. 22, 2015. featureCounts (v. 1.4.6-p3) [65] was used to compute gene-level read counts.

## Acknowledgments

We would like to thank Russell Fletcher and Diya Das for their constant feedback on the *scone* method and R package. We are grateful to Bioconductor core developers for the thorough code review and to all the users that contributed issue reports, bug fixes, and pull requests on GitHub.

---

**Supplementary Figure 1:**
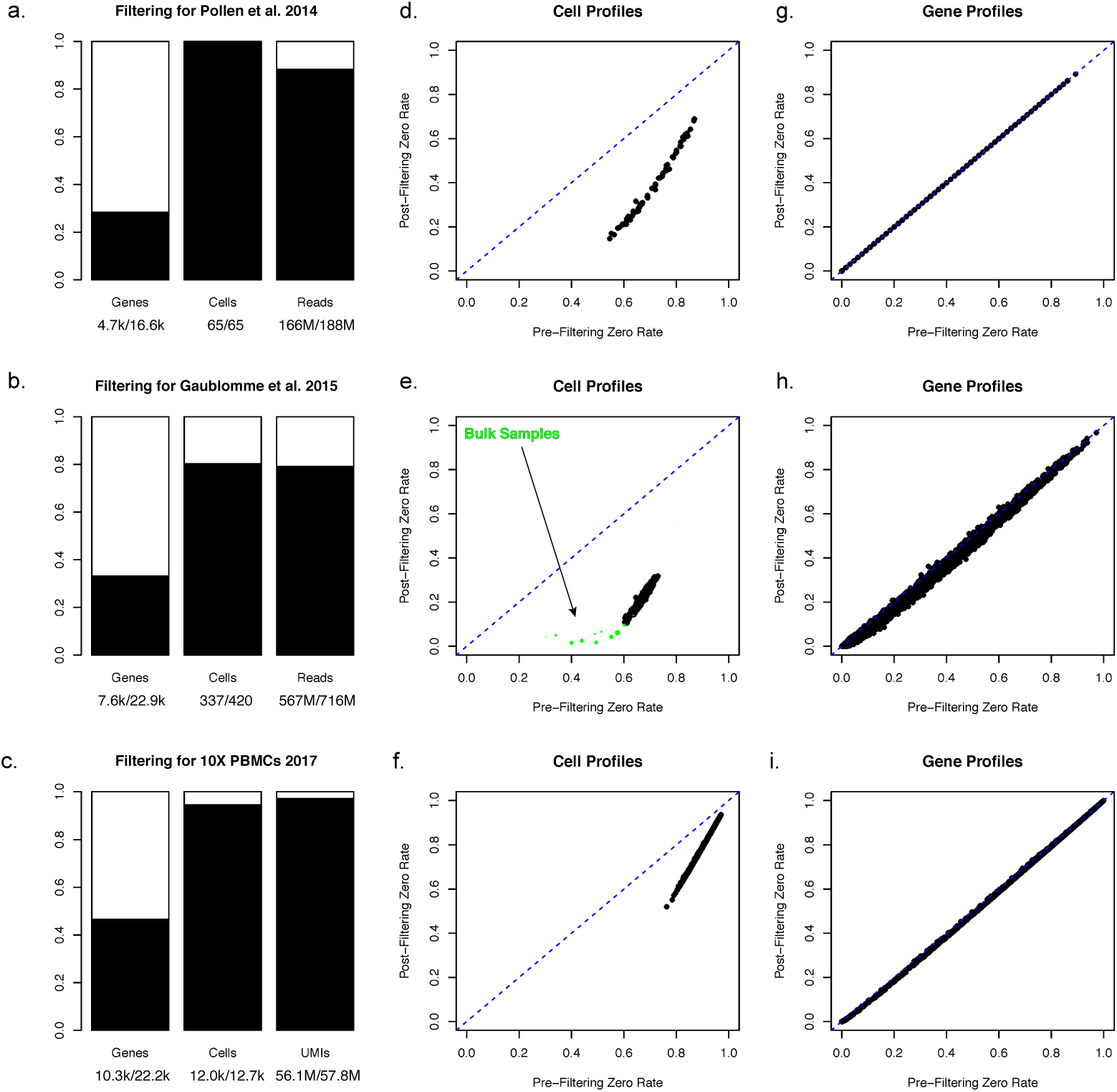
Data filtering for three scRNA-seq datasets [20, 22, 23]. (a-c) Boxplot representing (in black) proportions of genes, cells and counts preserved after sample and gene filtering. Note in (a) that no samples were filtered in Pollen et al. 2014. (d-f) Per-cell zero rates before and after data filtering. Datasets are presented in the same order as (a-c). In the case of Gaublomme et al. 2015, bulk RNA-seq samples from the same study were plotted in green along single-cell samples in (e). Point size corresponds to two-sided *p*-value of sample read count under log-normal model fit to single-cell samples passing filter. This is meant to highlight bulk samples with similar read coverage: other samples had very poor coverage and correspondingly high zero rates. (g-i) Per-gene zero rates before and after data filtering. Datasets are presented in the same order as (a-c).

**Supplementary Figure 2:**
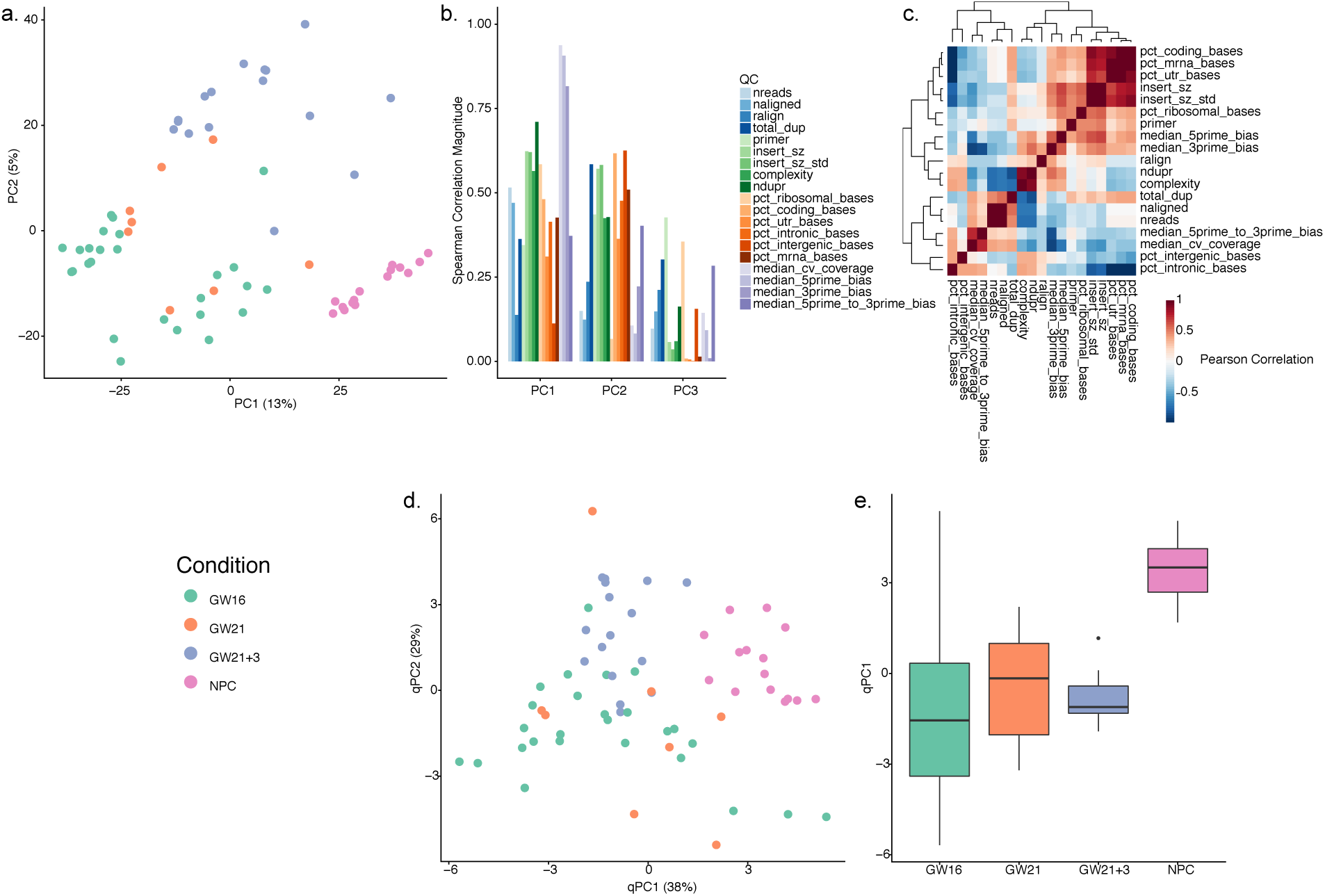
Exploratory data analysis of human cortex cells [22]. (a) PCA of the log-transformed, TC-normalized read count data using all genes passing quality filtering. Cells are color-coded by biological condition. Cells cluster partially by biological condition, with significant intra-condition heterogeneity. The design of this study is fully confounded (one batch per biological condition): batch adjustment is not advisable in this case, as it would remove the biological effects of interest. (b) Absolute Spearman correlation coefficient between the first three PCs of the expression data (as computed in (a)) and a set of QC measures (Table 1). (c) Heatmap of pairwise Pearson correlation coefficients between QC measures. (d) PCA of the QC measures for all cells in (a). Single-cell QC profiles cluster by biological condition, suggestive of technical confounding. (e) Boxplot of the first qPC, stratified by biological condition. QC measures differ significantly between NPCs and other biological conditions / batches.

**Supplementary Figure 3:**
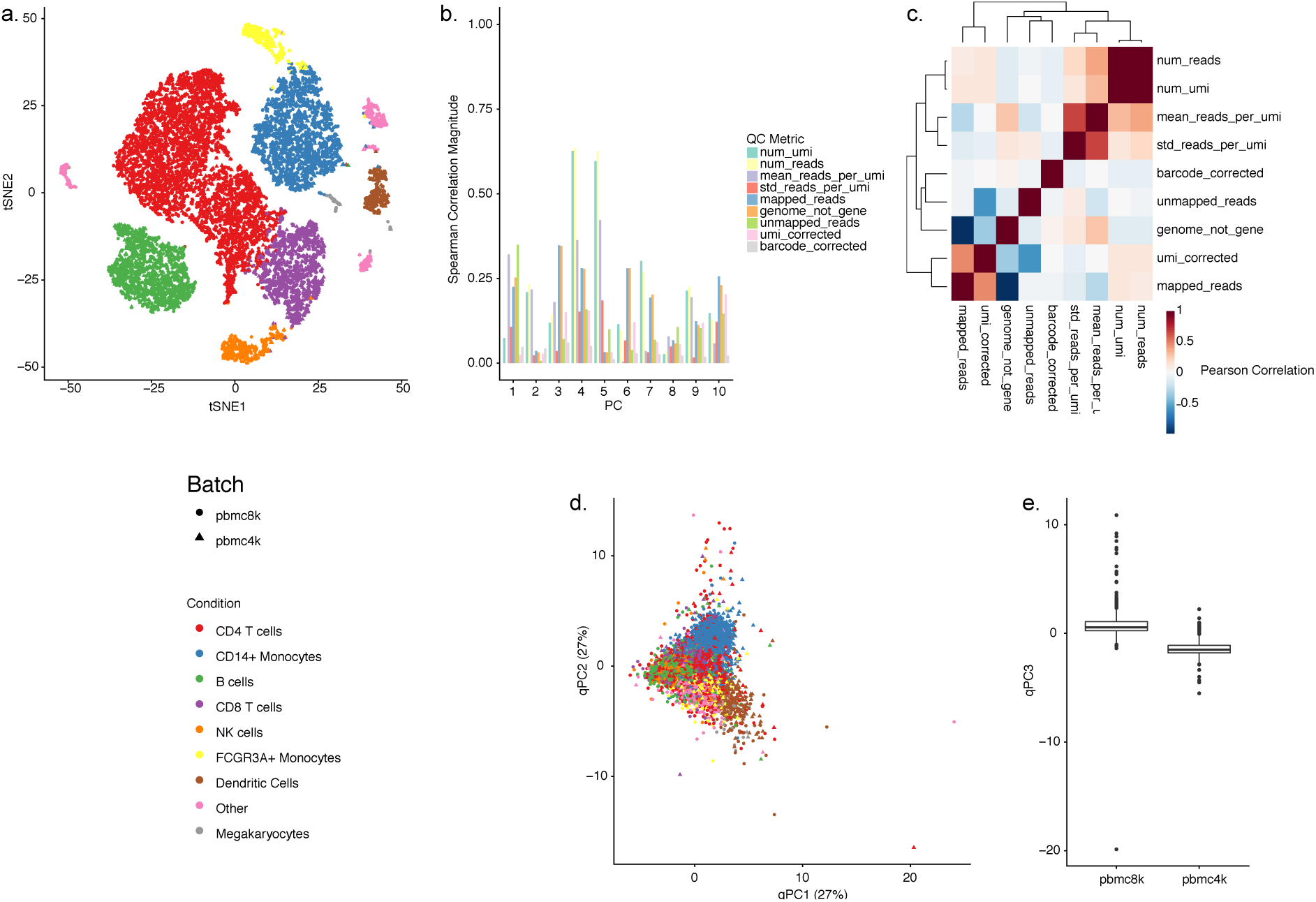
Exploratory data analysis of human peripheral blood mononuclear cells (PBMCs) [23]. (a) tSNE of the first 10 PCs of the log-transformed, TC-normalized UMI count data for all genes and cells passing quality filtering. Cells are color-coded by a Seurat-based manual annotation of major PBMC subtypes; shape represents the 10x batch. Samples from both batches (“pmbc4k" and a larger “pbmc8k") originated from the same “healthy" human donor. Cells clearly cluster by data-derived biological condition, one consequence of being clustered jointly in Seurat. (b) Absolute Spearman correlation coefficient between the first ten PCs of the expression data (as computed in (a)) and a set of QC measures (Table 2). (c) Heatmap of pairwise Pearson correlation coefficients between QC measures. (d) PCA of the QC measures for all cells in (a). Single-cell QC profiles partially cluster by data-derived biology (especially CD14+ monocytes), with no clear clustering by batch. (e) boxplot of the third qPC, stratified by batch. The third qPC is the qPC with the highest correlation with batch.

**Supplementary Figure 4:**
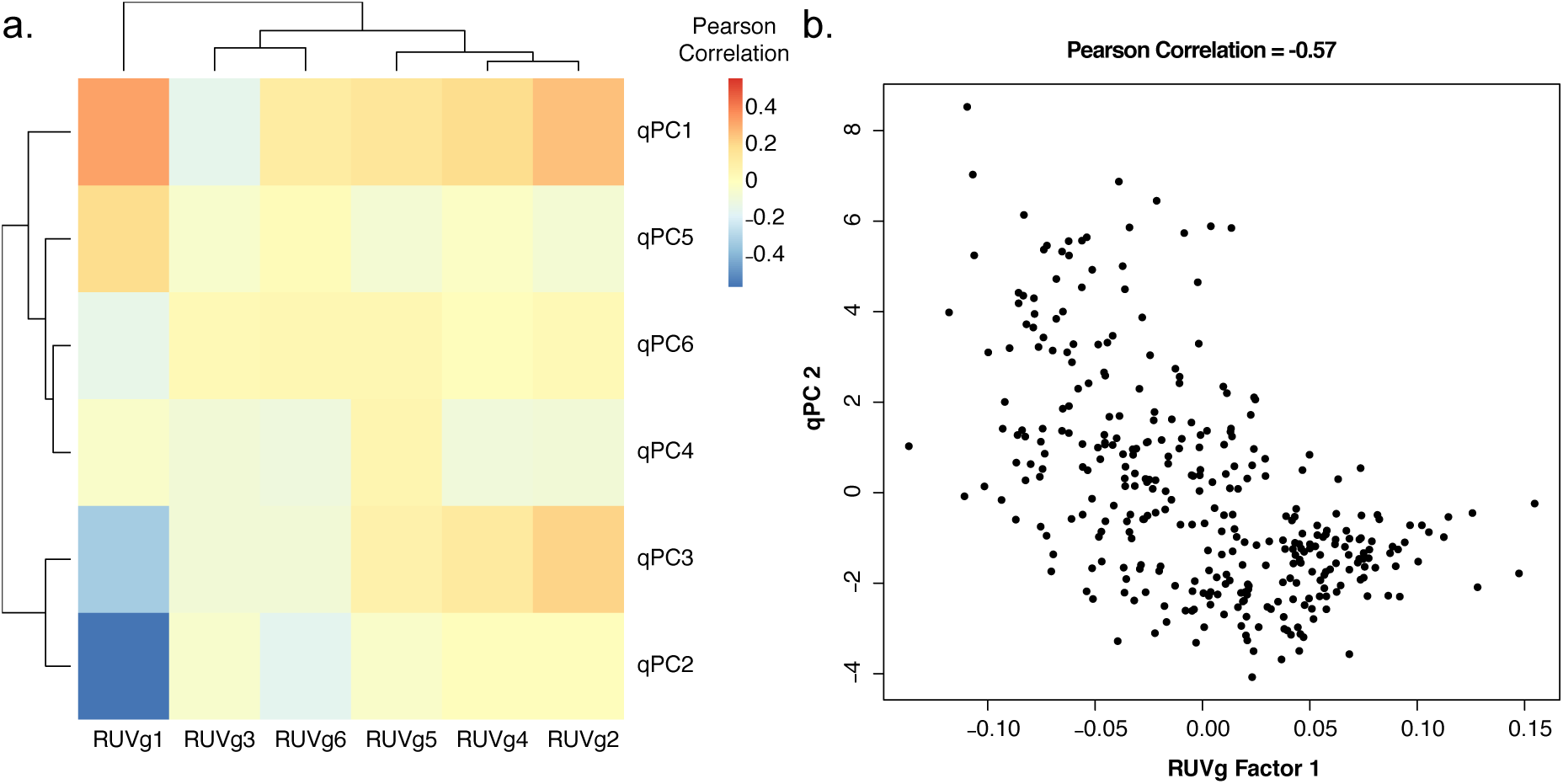
Factors of unwanted variation in the Gaublomme et al. dataset [20]. (a) Heatmap of Pearson correlation coefficients between RUVg-derived factors of unwanted variation [11] and qPCs. Row and column clustering is generated from the R hclust function with default parameters. (b) Scatter plot of one anti-correlated pair of RUVg factor and qPC, selected based on their high correlation magnitude displayed in (a).

**Supplementary Figure 5:**
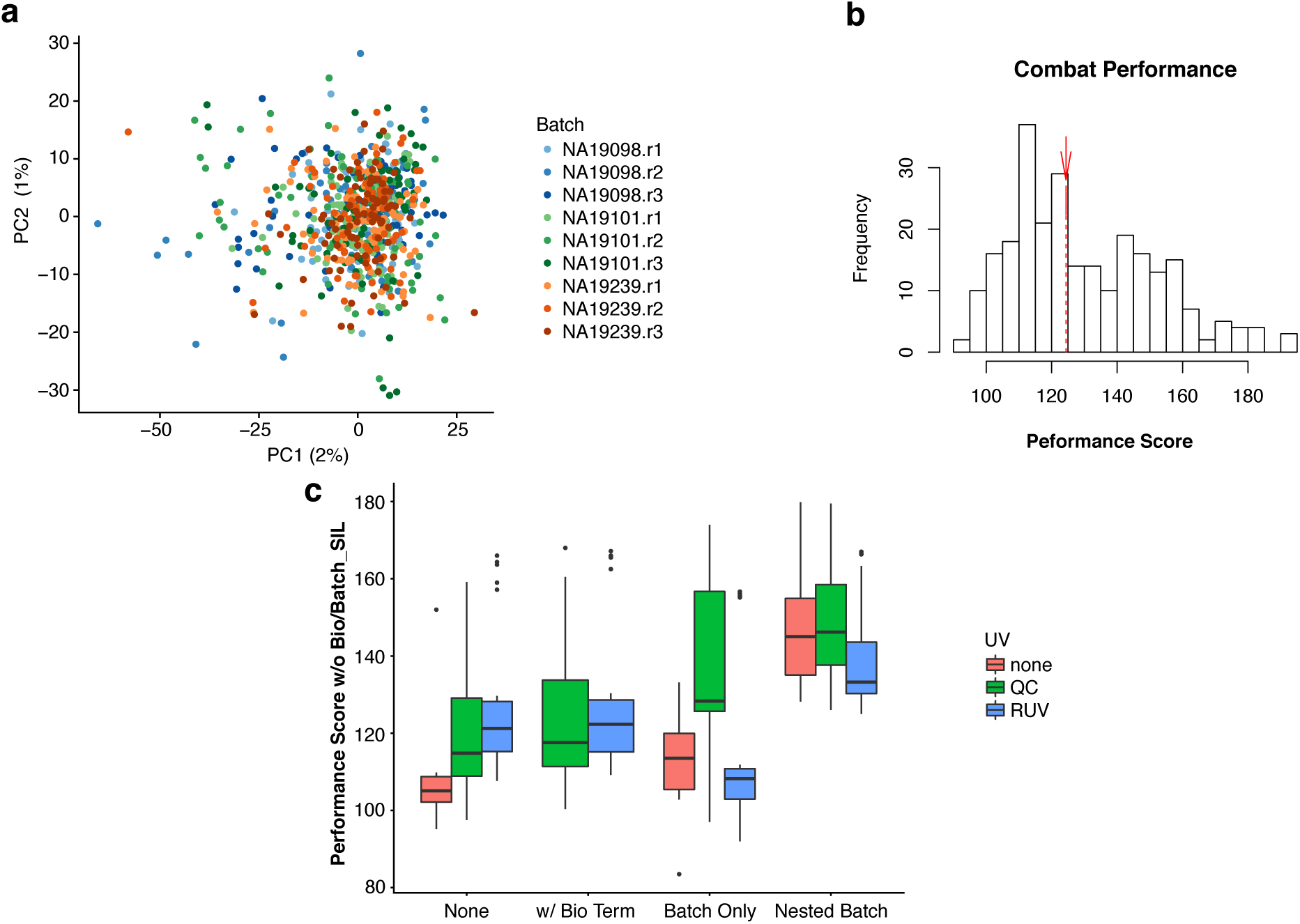
Batch adjustment for the Tung et al. dataset [32]. (a) PCA of ComBat [33] normalized data. Donor-specific effects are removed. (b) Histogram of *scone* performance scores recomputed to include ComBat (red arrow). (c) Boxplot of *scone* performance scores for various normalization procedures, excluding BIO_SIL and BATCH_SIL from the performance score calculation (see Methods).

